# RBL2 represses the transcriptional activity of Multicilin to inhibit multiciliogenesis

**DOI:** 10.1101/2023.08.04.551992

**Authors:** Erik J. Quiroz, Seongjae Kim, Lalit K. Gautam, Zea Borok, Christopher Kintner, Amy L. Ryan

**Author notes:** **Corresponding Author Contact Information**, Amy L. Ryan, PhD, Associate Professor: Anatomy and Cell Biology, Associate Director: Center for Gene Therapy, BSB, 1-400 Core, University of Iowa 51 Newton Road, Iowa City, Iowa, 52242. These authors contributed equally to this manuscript (Erik J. Quiroz, Seongjae Kim, Amy L. Ryan).

## Abstract

A core pathophysiologic feature underlying many respiratory diseases is multiciliated cell dysfunction, leading to inadequate mucociliary clearance. Due to the prevalence and highly variable etiology of mucociliary dysfunction in respiratory diseases, it is critical to understand the mechanisms controlling multiciliogenesis that may be targeted to restore functional mucociliary clearance. Multicilin, in a complex with E2F4, is necessary and sufficient to drive multiciliogenesis in airway epithelia, however this does not apply to all cell types, nor does it occur evenly across all cells in the same cell population. In this study we further investigated how co-factors regulate the ability of Multicilin to drive multiciliogenesis. Combining data in mouse embryonic fibroblasts and human bronchial epithelial cells, we identify RBL2 as a repressor of the transcriptional activity of Multicilin. Knockdown of RBL2 in submerged cultures or phosphorylation of RBL2 in response to apical air exposure, in the presence of Multicilin, allows multiciliogenesis to progress. These data demonstrate a dynamic interaction between RBL2 and Multicilin that regulates the capacity of cells to differentiate and multiciliate. Identification of this mechanism has important implications for facilitating MCC differentiation in diseases with impaired mucociliary clearance.

## Introduction

Multiciliated cells (MCCs) are key drivers of directional ciliary flow along epithelia in many tissues including the airways, reproductive tract, and brain^1–6^. In the airways, MCCs play an integral role in the lung’s primary innate defense system, providing both a physical barrier from the environment and mechanical force that drives mucus, pathogens, and foreign debris out of the lungs in a process called mucociliary clearance^7^. Due to their central role in maintaining airway homeostasis, dysregulated MCC function is a critical pathophysiologic feature underlying many respiratory diseases, including chronic obstructive pulmonary disease (COPD) and cystic fibrosis (CF)^8, 9^. Primary ciliary dyskinesia (PCD)^10^, a genetic disease arising from mutations in cilia related genes, results in non-motile, dyskinetic, or absent cilia, and symptomatically recapitulates secondary ciliary dyskinesia, characterized as the disruption of ciliary function from disease or injury^11, 12, 13, 14^. Repeated injury, characteristic of chronic lung disease, can lead to airway remodeling and changes in the cellular composition of airway epithelium, resulting in a decreased number of MCCs covering the airway surface^8^. Due the prevalence of MCC dysfunction in respiratory diseases, it is therefore critical to understand the mechanisms controlling multiciliogenesis and how these may be targeted to restore functional mucociliary clearance.

To accomplish their function, each airway MCC must generate ∼150 motile cilia^15^. Cilia are microtubule-based organelles anchored to the plasma membrane by a basal body, formed from a mature parent centriole during cellular quiescence^15^. Most cell types extend a single cilium because centriole number is tightly controlled, with centriole duplication occurring only once per cell cycle^16^. By contrast, differentiating MCCs generate hundreds of centrioles as post-mitotic cells, thus engaging novel centriole biogenesis mechanisms outside the cell cycle and in a manner capable of significantly increasing centriole number. This process of massive centriole amplification is mediated by a MCC-specific transcriptional program that induces expression of the centriolar proteins that allow for centriolar duplication, which normally occurs only during S phase, as well proteins that form a structure unique to MCCs known as the deuterosome^17–23^.

Geminin family proteins, GEMC1 and Multicilin, encoded by the gene *MCIDAS*, have been identified as master transcriptional regulators of multiciliogenesis, controlling both the induction of deuterosome-mediated centriole biogenesis and activating the gene regulatory network necessary to produce motile cilia^4, 24–27^. GEMC1 and Multicilin share similar sequence homology, with a conserved coiled-coil domain and carboxy terminus; however, GEMC1 has been shown to help specify MCC fate, while Multicilin is sufficient to induce multiciliogenesis and can bypass GEMC1^28^. Upstream of GEMC1 and Multicilin, inhibition of NOTCH signaling is critical in regulating multiciliogenesis; a direct link between NOTCH-mediated cell fate determination and induction of Multicilin remains unknown^28^. During multiciliogenesis, Multicilin acts transcriptionally in a complex with cell cycle transcription factors E2F4 or E2F5, along with dimerization partner DP-1, activating the expression of centriole biogenesis genes and key downstream transcription factors, including c-MYB, TP73, and FOXJ1^24^. Paradoxically, this activation occurs despite the well-known role of the E2F4 and 5 in post-mitotic cells as repressors of gene expression during cell cycle progression, by acting in a complex with the retinoblastoma (RB) family of co-repressors, and the multi-vulval class B (DREAM) complex^29^.

While Multicilin is known to be sufficient to drive multiciliogenesis in certain cell types^30–32^, it has become apparent that this does not apply to all cell types, nor occur evenly across all cells in the same cell population. In our prior study using mouse embryonic fibroblasts (MEFs), expression of Multicilin alone was insufficient and could only stimulate multiciliogenesis after serum starvation and when co-expressed with E2f4 modified by replacing the C-terminus with a VP16 transactivator, indicating that the Multicilin requires a specific cellular context in order to promote multicilogenesis^33^. In this report we further investigate how co-factors regulate the ability of Multicilin to drive multiciliogenesis. We identify RBL2 as a repressor of the transcriptional activity of Multicilin and demonstrate its dynamic interaction during differentiation of human airway epithelial cells at the air-liquid interface (ALI). Identification of this mechanism has important implications for facilitating MCC differentiation in diseases where mucociliary clearance is impeded.

## Results

### E2f4 C-terminus substitution with VP16 attenuates Rbl2-Multicilin interaction in MEF

In their canonical roles, the E2F4 pocket protein binding domain (PPBD) is known to bind co-repressors to inhibit cell cycle gene expression, while the transactivating domain (TAD) of E2F family proteins associates with epigenetic modifiers to activate transcription (**Supplementary Figure S1a**)^34^. We have previously reported that overexpression of Multicilin was not sufficient to induce multiciliogenesis effectively in MEFs^33^. This block could be overcome by overexpressing Multicilin along with a modified form of E2f4 where both the PPBD and TAD domains were replaced with a viral transactivation domain VP16 (E2f4ΔCT-VP16), as evidenced by robust centriole amplification and increased expression of multiciliogenesis-related proteins (**Supplementary Figure S1b,c**)^33^. To determine whether the proteins associated with the Multicilin complex change when cells express Multicilin along with wildtype E2f4 (limited multiciliogenesis) compared to E2f4ΔCT-VP16 (extensive multiciliogenesis) we performed proteomics.

Adenoviral vectors were used to express Multicilin tagged at the N-terminus with 3xFLAG (FLAG-MCI, **Fig. 1a**), FLAG-MCI along with wildtype E2f4 (FLAG-MCI+E2f4, **Fig. 1b**), or FLAG-MCI along with E2f4ΔCT-VP16 (FLAG-MCI+E2f4VP16, **Fig. 1c**). MEFs were transduced, and immunoprecipitation performed using FLAG to co-purify proteins complexed to Multicilin. The bound proteins to FLAG-MCI were eluted and silver staining performed to identify the MCI-3xFLAG (**Fig. 1e**). Mass spectrometry analysis identified multiple proteins with similar abundance between FLAG-MCI and FLAG-MCI+E2f4VP16, with 8 of the 10 most abundant proteins being present in both conditions. These proteins included known Multicilin binding partners, Tfdp-1, Gmnn, E2f4, and E2f5, as well as ribosomal proteins whose interaction with Multicilin has not previously been described: Rps27, Rps27l and Rps14 (**Fig. 1d)**. A complete list of the proteins complexed with Multicilin is included in **Supplementary Table S1**. The other 2 of the 10 most abundant proteins bound to FLAG-MCI were 26S protease regulatory subunit 7 (Psmc2) and Retinoblastoma-like protein 2 (Rbl2). Notably, Rbl2 had the largest decrease in relative abundance comparing FLAG-MCI to E2f4VP16 of all proteins detected in both samples, with a ratio of 9.2:1.

**Figure 1:**
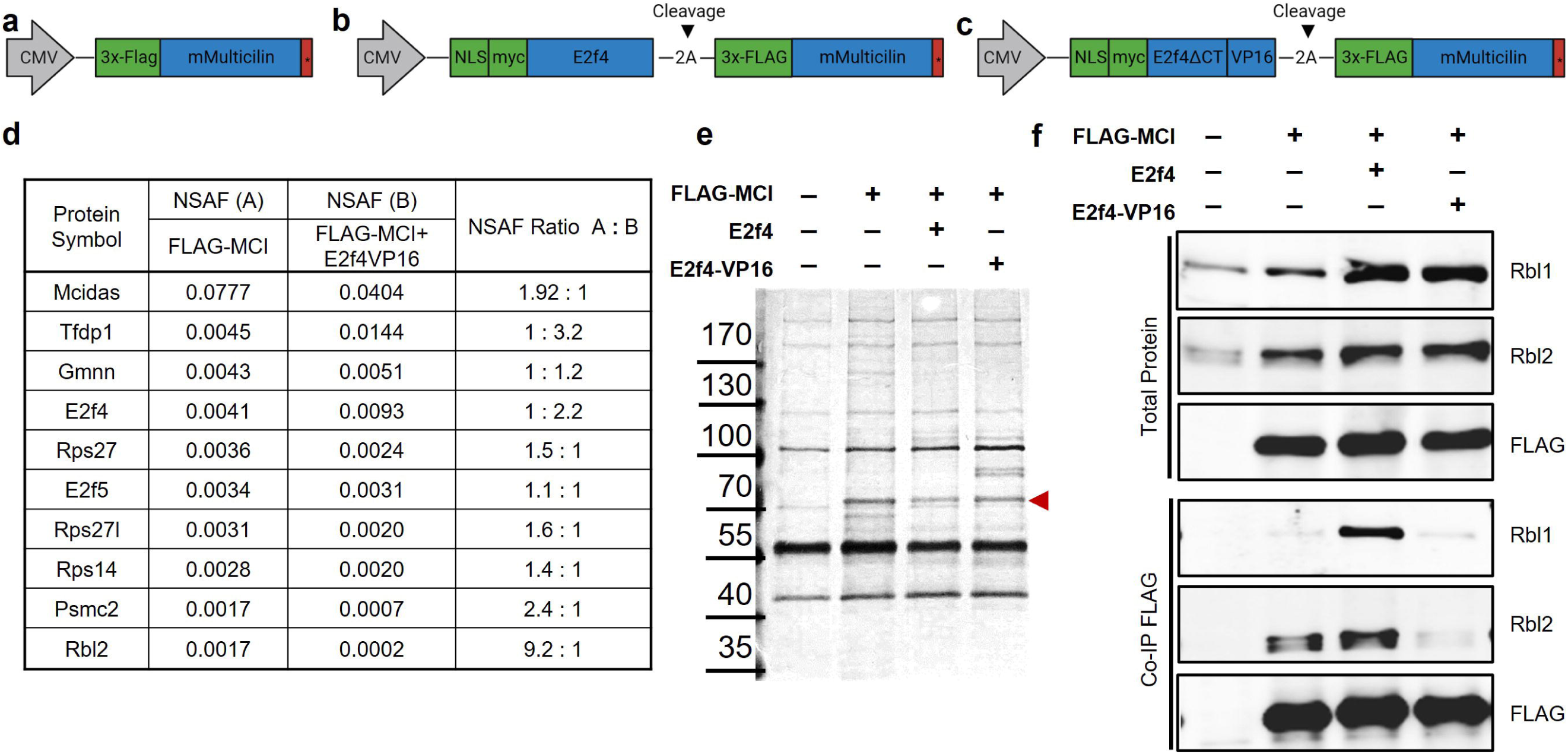
Multicilin physically interacts with Rbl2 in mouse embryonic fibroblasts. (**a-c**) Schematics of the adenoviral constructs for FLAG-MCI (**a**), FLAG-MCI+E2f4 (**b**), and FLAG-MCI+E2f4VP16 (**c**). (**d**) Normalized spectral abundance factor (NSAF) for the top 10 most abundant proteins co-immunoprecipitating with FLAG-MCI relative to FLAG-MCI+E2f4VP16 (see **Supplemental Table 1**). (**e**) Representative silver stain of total protein lysate, the red arrowhead indicates 3xFLAG-MCI. (**f**) Representative Western blot analysis for Rbl1, Rbl2, and FLAG for total protein (top) and co-immunoprecipitations (Co-IP, bottom) with FLAG from non-infected mouse embryonic fibroblasts (MEFs) and MEFs infected with Ad5 FLAG-MCI, FLAG-MCI+E2f4 and FLAG-MCI+E2f4VP16.

Immunoblotting analysis detected a strong interaction between Multicilin and Rbl2 in MEFs expressing either FLAG-MCI or FLAG-MCI+E2f4; however the Rbl2-Multicilin interaction was abolished in MEFs expressing FLAG-MCI+E2f4VP16 (**Fig. 1f**). Similarly, an interaction with Rbl1 was observed in MEFs expressing FLAG-MCI in the presence of E2f4, but not E2f4VP16 (**Fig. 1f**). The retinoblastoma family of proteins, including Rb, Rbl1 and Rbl2, have been well characterized in their roles as cell cycle inhibitors through interactions with the E2f family of transcription factors^35, 36^. The Rbl2-E2f4 complexes have been identified as the predominant repressors of cell cycle genes during quiescence^35, 36^. Taken together, the canonical role of Rbl2 as a co-repressor of E2f4 and our observation that Rbl2 complexes strongly with Multicilin in the presence of E2f4 relative to E2f4ΔCT-VP16, suggest that Rbl2 may regulate the capacity of the Multicilin transcriptional complex to activate MCC gene expression.

### Rbl2 represses the ability of Multicilin to drive ectopic multiciliogenesis in MEFs

To further investigate a regulatory role of Rbl2 in the Multicilin-E2f4 transcriptional complex, we tested siRNAs for *Rbl2* knockdown in MEFs (siRBL2#1-3, **Supplementary Table S2**), identifying siRbl2#1 (siRbl2) as most efficient at reducing *Rbl2* gene expression, with a reduction of 1.42±0.44 log2 fold change compared to a non-targeting control (siCTL) (**Supplemental Figure S2a**). An siRNA targeting *Rbl1* (siRbl1) was also tested and compared as an additional control. Successful knockdown was validated at the protein level by Western blot (**Fig. 2a, Supplementary Figure S2b-c)**.

**Figure 2:**
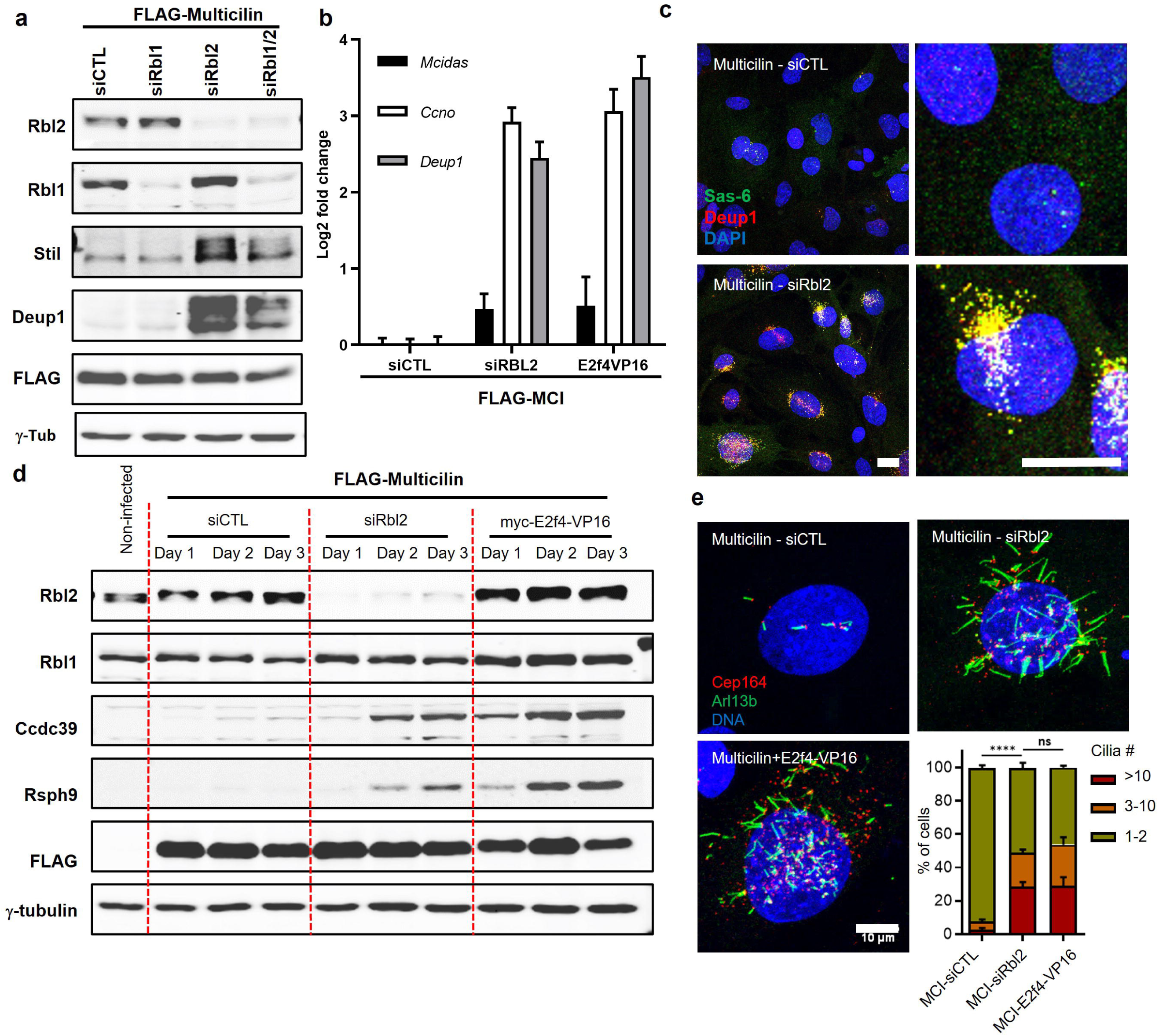
Rbl2 depletion increases the transcriptional activity of Multicilin stimulating multiciliogenesis in fibroblasts. (**a**) Representative Western blot for Rbl2, Rbl1 and centriole biogenesis proteins Stil and Deup1, comparing FLAG-Multicilin in the presence of a non-targeting siRNA (siCTL) or siRNA targeting *Rbl1* (siRBL1), *Rbl2* (siRbl2), or both siRBL1 and siRBL2 (siRBL1/2). (**b**) qRT-PCR for MCC genes *Mcidas*, *Ccno* and *Deup1* comparing FLAG-Multicilin with either siRBL2 or E2f4VP16 relative to siCTL (Data expressed as mean±SEM., N=2 experimental replicates, normalized to Gapdh). (**c**) Confocal images comparing FLAG-Multicilin in the presence of either siCTL or siRBL2 stained for centriole biogenesis markers Deup1 (red), Sas-6 (green) and nuclei are counterstained with DAPI (blue). Scale bars = 20 µm. (**d)** Representative Western blot comparing FLAG-Multicilin in the presence siCTL, siRBL2, or E2f4VP16 for Rbl2, Rbl1, FLAG, axonemal proteins CCDC39 and RSPH9 and γ-tubulin (γ-tub, loading control). (**e**) Confocal images for cilia marker Arl3b (green) and basal body marker Cep164 (red) nuclei are counterstained with DAPI (blue). (**f**) Cilia are quantified per cell, data represents mean±SD, N=3, **** P<0.0001, unpaired t-test. Scale bar = 10 µm.

Multicilin-expressing, Rbl2-depleted MEFs (FLAG-MCI+siRbl2) showed a marked increase in the transcriptional activation of core multiciliogenesis genes *Ccn*o, *Deup1 and Mcidas* relative to control siRNA MEFs (FLAG-MCI+siCTL), reaching levels comparable to those in Multicilin-expressing MEFs with E2f4VP16 (FLAG-MCI-E2f4VP16) **(Fig. 2b)**. Similar increases were observed at the protein level, since a depletion of *Rbl2*, but not *Rbl1*, in Multicilin-expressing MEFs significantly induced the expression of centriolar assembly proteins, Stil and Deup1 (**Fig. 2a),** as well as ciliary axonemal protein, Ccdc39 (**Supplemental Figure S2b**). In Multicilin-expressing MEFs with siRbl2, robust centriole amplification was observed as shown in representative confocal images showing Deup1 and Sas6, markers of centriole amplification via the deuterosome pathway (**Fig. 2c, Supplemental Figure S2d)**. Immunoblotting confirmed the presence of axonemal proteins, Ccdc39 and Rsph9, in both siRbl2 and E2f4VP16, but not siCTL, indicating sufficient activation of late stage multiciliogenesis by Multicilin only in the absence of Multicilin interaction with Rbl2 (**Fig. 2d**). The formation of cilia was validated by immunostaining with cilia marker, Arl3b, and basal body marker, Cep164, revealing that Multicilin induction stimulated cilia differentiation Rbl2-depleted and E2f4VP16 expressing, but not in control siRNA-transfected MEFs (**Fig. 2e**).

To further evaluate the ability of Rbl2 to regulate Multicilin-dependent activation of multiciliogenesis, we performed bulk RNA-sequencing to determine Multicilin-induced transcriptional changes comparing Multicilin induced MEFS with siCTL, siRbl2 or E2f4VP16 to non-infected MEFs (**Supplementary Table S3)**. Multicilin-induction with E2f4VP16 induced the most dramatic changes in overall gene expression compared to the non-infected MEFs, with similar trends occurring in the presence of siRbl2 and with siCTL having less substantial transcriptional changes (**Supplemental Figure S3**). Evaluating a panel of MCC-related genes^37^, both E2f4VP16 and siRbl2 induce comparable transcriptional changes, while siCTL more closely reflects non-infected MEFs (**Fig 3a, Supplemental Figure S4)**. Gene Ontogeny (GO) analysis of differentially expressed genes (DEGs) comparing experimental conditions and the non-infected controls (adj p>0.05, |FC|>2) highlighted significant enrichment in Multicilin-induced MEFs, in the presence of either E2f4VP16 or siRbl2, for biological processes (GO:BPs) such as cilium organization, cilium movement, and protein localization to cilium as well cellular components (GO:CCs) cilium, centriole, manchette and dynein axonemal particle (**Fig. 3b**). Multicilin-induction with siCTL had a significant enrichment in GO:CC deuterosome and GO:BPs cilium organization and multiciliated differentiation; however, the most significantly enriched GO:BPs were immune system process and response to interferon-beta (**Fig. 3b**). Similar numbers of DEGs from these GO terms were also noted in the E2f4VP16 and siRbl2 MEFs, however the greater total number of DEGs decreased the enrichment significance for these GO terms (**Supplementary Table S3)**. Comparing common DEGs between samples, Multicilin-induced MEFs with E2f4VP16 have higher MCC gene expression compared to Rbl2-depleted MEFs (**Fig 3c**), however both are significantly increased compared to siCTL (**Fig 3d-e**). The Venn diagram shown in **Figure 3f** compares all DEGs, with >2-fold changes in expression, across all experimental conditions relative to non-infected controls. We identified 492 DEGs significantly upregulated specifically in the Multicilin-induced and Rbl2-depleted or E2f4VP16 expressing MEFs, but not in the Multicilin-induced siCTL MEFs (**Fig. 3g**). Of these 492 genes, 225 are known MCC-associated genes (**Fig. 3g)**^37^. This data supports a regulatory role for the suppression of Rbl2 in enabling Multicilin to activate transcriptional pathways to induce massive centriole biogenesis and form multiple motile cilia.

**Figure 3.**
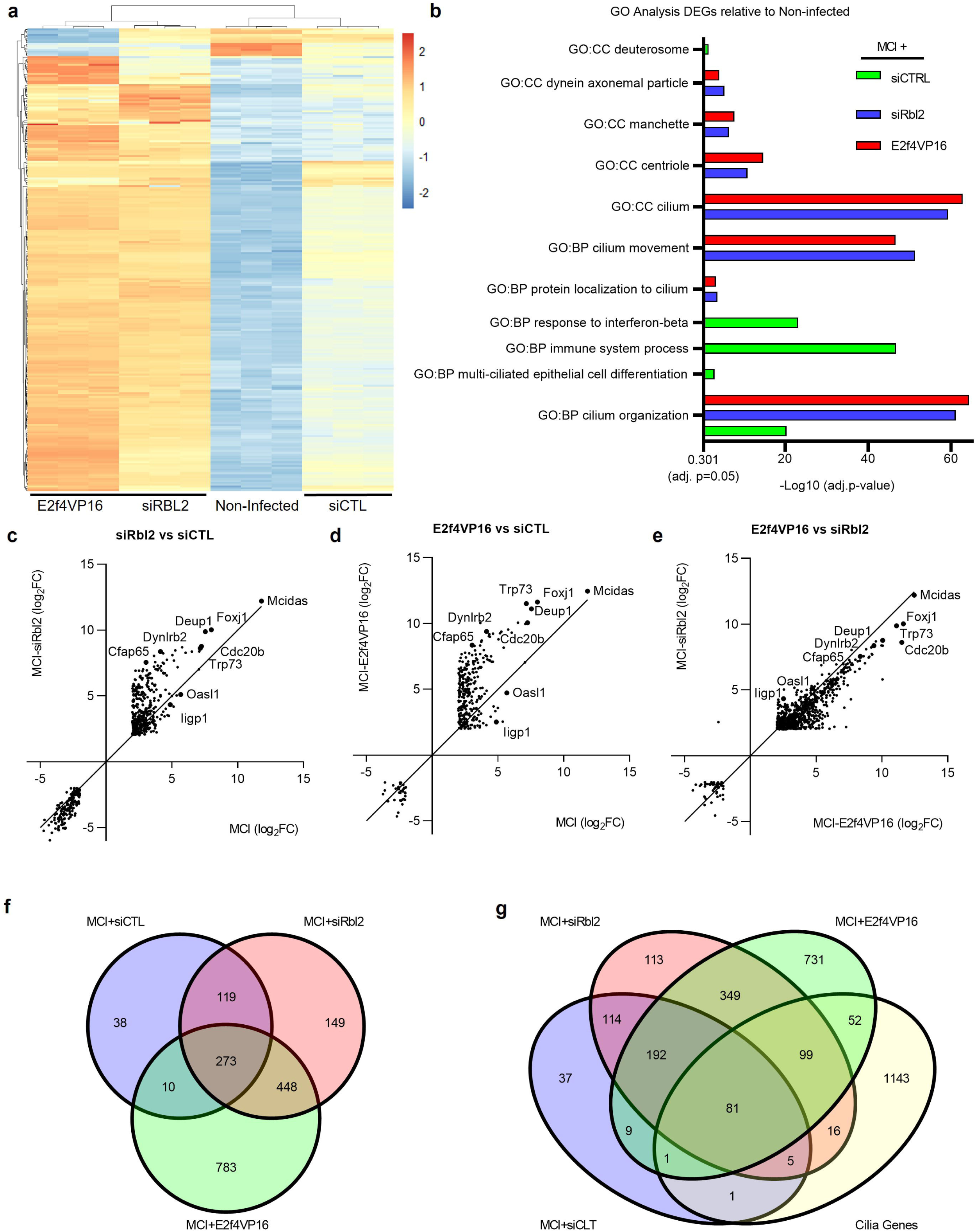
Transcriptome profiling confirms induction of multiciliogenesis in the presence of siRbl2 or E2f4VP16 compared to Multicilin alone. (**a**) Unsupervised heatmap of RNA-seq data for MCC-associated genes comparing non-infected MEFs and FLAG-MCI MEFs with either siCTL, siRBL2 or E2f4VP16. (**b**) Gene ontology (GO) analysis of differentially expressed genes (DEGs) (**Supplemental Table 3)** comparing non-infected MEFs and FLAG-MCI MEFs with either siCTL, siRBL2 or E2f4VP16. GO terms include biological process (GO:BP) and cellular components (GO:CC). (**c-e**) Scatter plots comparing significant DEGs in FLAG-MCI MEFs (relative to uninfected, absolute log2 fold change>2, adj. p<0.5) for siRBL2 vs siCTL **(c)**, E2f4VP16 vs siCTL (**d**), and siRBL2 vs E2f4VP16 (**e**). (**f-g**) Venn diagrams representing significantly increased DEGs in FLAG-MCI relative to uninfected controls (log2 fold change>2, adj. p<0.5, **Supplemental Table S5**) comparing siRBL2, siCTL and E2f4VP16 (**f**) and compared to known MCC-specific genes^37^ (**g**). N=3 experimental replicates for all RNAseq data.

### RBL2 expression does not change the extent of multiciliogenesis from HBECs

Since knockdown of *Rbl2* allowed Multicilin to induce ectopic multiciliogenesis in MEFs, a cell type that is not innately capable of multiciliogenesis, we investigated whether *RBL2* knockdown in human MCC progenitors can potentiate multiciliogenesis. To accomplish this, we compared the differentiation of human bronchial epithelial cells (HBECs) at the air-liquid interface (ALI) with or without *RBL2* knockdown (**Fig. 4**). qRT-PCR analysis after 14 days of ALI differentiation confirmed that shRBL2 (shRBL2#1 and #2) significantly decreased expression of *RBL2* when compared to non-targeting controls (shNTC#1 and #2), by 2.37±0.63 and 1.30±0.19 log2 fold change, respectively (**Fig. 4a**). The reduction in *RBL2* expression resulted in a modest increase in the expression of early MCC differentiation genes including *SAS6, DEUP1* and *TP73* (**Fig. 4b**), however the expression of genes associated with differentiated human airway epithelium, including goblet (*MUC5AC)*, club (*CC10)*, and ciliated (*FOXJ1) cells* was unchanged (**Fig. 4c**). This suggests that the proportion of MCCs generated in the presence of *RBL2* knockdown was unchanged. Immunoblotting confirmed knockdown of RBL2 at the protein level alongside increased expression of TP73, while more mature MCC markers, FOXJ1 and axonemal protein CCDC39, remained unchanged (**Fig. 4d-e**). Immunostaining for cilia maker acetylated α-tubulin (ATUB) was used to evaluate the efficiency of MCC differentiation by quantifying the area of ciliated cell coverage on the apical surface at ALI day 14, while live video analysis was used to measure cilia beat frequency (CBF). No significant changes in total ciliated cell coverage, nor in CBF were detected in RBL2 knockdowns compared to controls (**Fig. 4g-h**). These data show that ectopic reduction of RBL2 expression does not change MCC maturation and function and suggests that RBL2 activity may already be endogenously regulated during the early stages of HBEC differentiation at the ALI.

**Figure 4:**
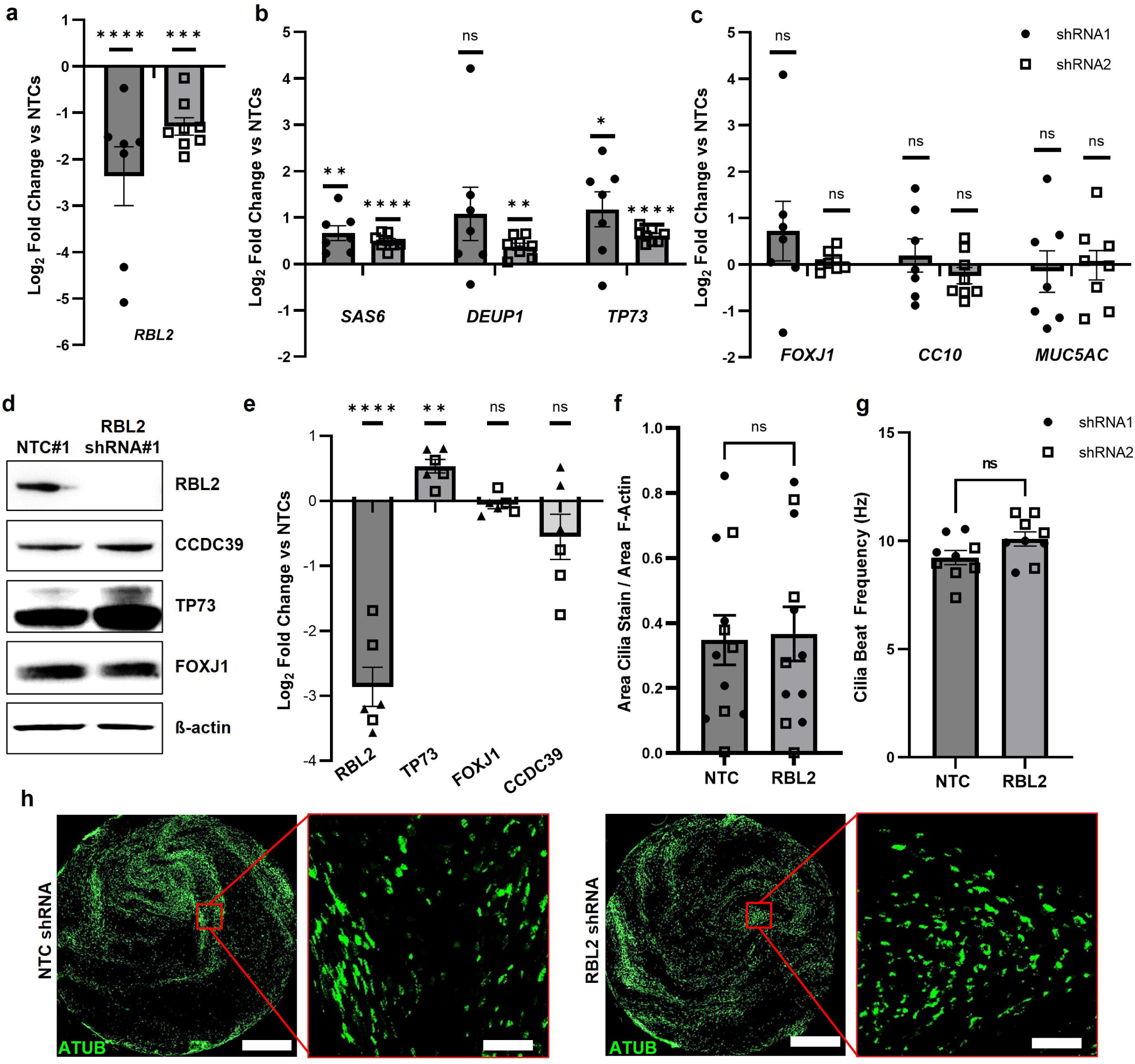
RBL2 knockdown does not impact the efficiency of MCC differentiation in HBEC. qRT-PCR comparing HBECs treated with shRNA-RBL2 (shRNA#1 (filled circles), shRNA#2 (open squares)) or a non-targeting control (NTC) at Day 14 if ALI culture for *RBL2* (**a**), early MCC markers *SAS6*, *Deup1* and *TP73* (**b**) and differentiated epithelial cell markers *FOXJ1* (MCC), *CC10* (club), and *Muc5AC* (goblet) (**c**). Data is corrected for *RPLP0* and normalized to NTC shRNA, N=4-5 biological replicates each with n=1-2 experimental replicates paired for each shRNA. (**d**) Representative Western blot for shRNA#1 and NTC#1 at Day 14 ALI for RBL2 and MCC proteins CCDC39, TP73 and FOXJ1. **(e**) Quantification of protein expression from N=3 biological replicates for each shRNA. (**f**) Quantification of cilia coverage at D14 ALI expressed as a ratio of area of total acetylated α-tubulin (ATUB, cilia) over total growth area (area F-Actin), N=5-6 biological replicates each n=1-2 experimental replicates. (**g)** Analysis of cilia beat frequency (CBF) at D14 ALI, N=4-5 biological replicates. (**h**) Representative IF images of ATUB (green), scale bars represent 1 mm and 100 µm. Data is expressed as mean±SEM and compared using two-tailed paired t-tests (a-c &e) ratio-paired t-test (f) paired t-test (g) and *P<0.05, ** P<0.01, *** P<0.001, **** P<0.0001.

### ALI culture phosphorylates RBL2, allowing Multicilin to drive multiciliogenesis in HBECs

To determine whether RBL2 interactions with Multicilin are indeed altered during the early phase of ALI differentiation of HBECs, we evaluated RBL2-Multicilin interactions comparing ALI to submerged cultures in differentiation media. Doxycycline (DOX) induction of lentiviral-transduced FLAG-tagged Multicilin in HBECs was performed over a 4-day period prior to sample collection (**Fig. 5a, Supplementary Figure S5a-b**). Immunoprecipitation of FLAG-tagged Multicilin and immunoblotting for RBL2 revealed a higher amount of RBL2 complexed with Multicilin in the submerged cultures (**Fig. 5a**, Sub D0 and Sub D4), compared to ALI cultures (**Fig. 5a**, ALI D4 and ALI D21). Total RBL2 remained consistent in all conditions (**Fig. 5a, Supplementary Figure S5b**). These data demonstrate that ALI culture attenuates the direct interaction between RBL2 and Multicilin in HBECs, which in turn potentiates Multicilin transcriptional activity to drive MCC differentiation.

**Figure 5:**
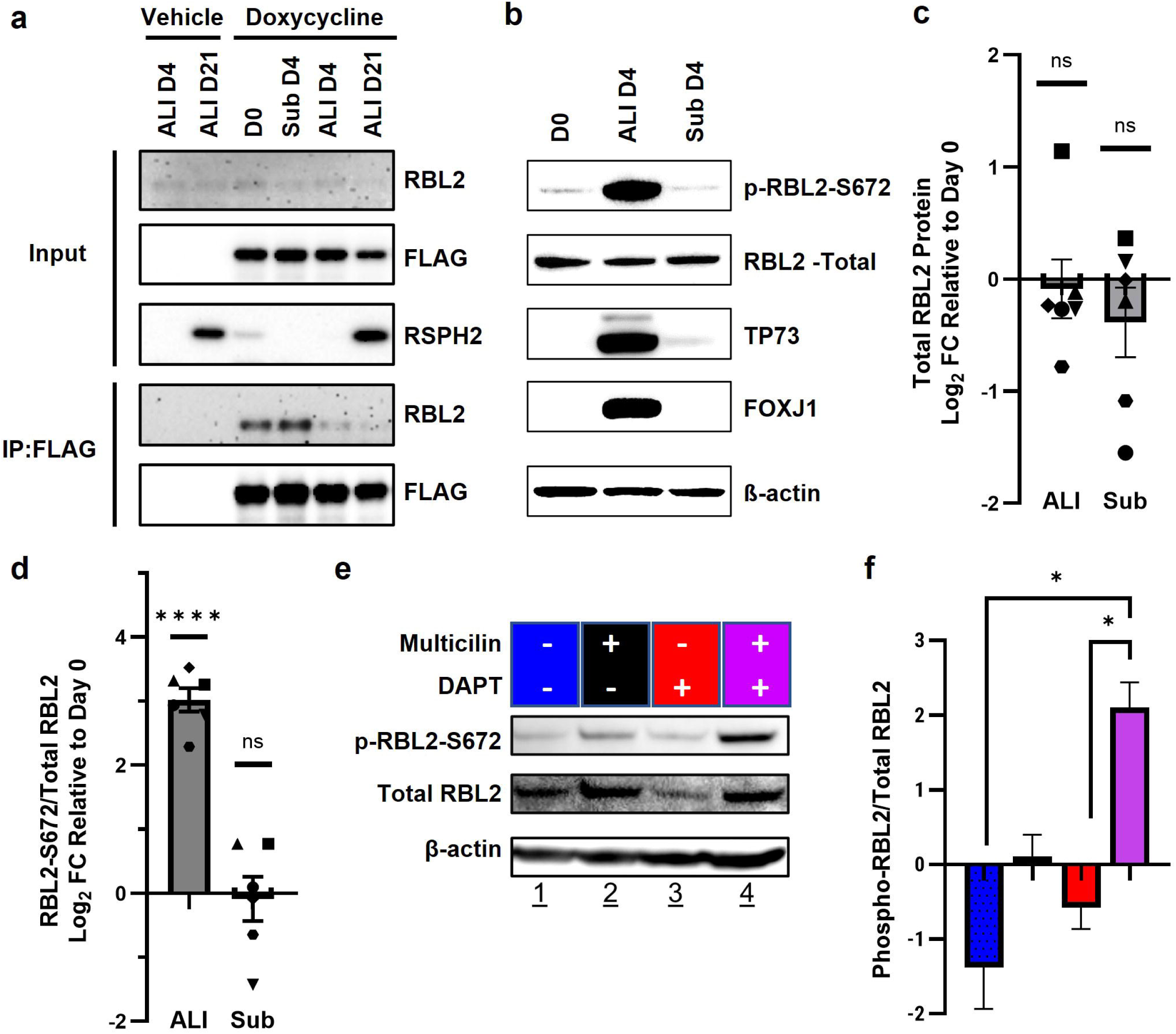
Air-liquid interface attenuates the interaction between RBL2-Multicilin. (**a**) Representative Western blot for total protein and co-IP for FLAG in FLAG-MCI induced HBECs either submerged (Sub) or ALI culture. (**b**) Representative Western Blot for total RBL2, phosphorylated serine 672 RBL2 (RBL2 S672) and MCC proteins TP73 and FOXJ1 at D0, D4 (Sub) and ALI D4. Protein quantification for total-RBL2 (**c**) and RBL2 S672:total RBL2 (**d**), N=6 biological replicates. (**e**) Representative Western blot for Multicilin-induced HBECs at Sub D4 in the presence of 5 µM DAPT or vehicle. (**f**) Protein quantification for DAPT treatment from N=3, from 1-2 biological replicates and 1-2 experimental replicates. Data represents mean±SEM and is compared using a paired t-test (c-d) and 2-way ANOVA and post-hoc Tukey’s multiple comparisons test (f).

Cyclin dependent kinases are known to phosphorylate the RB family of proteins, interrupting their interaction with E2F family transcription factors to regulate gene expression and cell cycle progression^38, 39^. Canonically, RBL1/2 interactions with E2F4/5 actively repress cell cycle genes^36^. The ability of Multicilin to co-opt E2F4 to induce centriole biogenesis in cells not actively dividing, suggests a mechanism whereby RBL2 may be phosphorylated and inactivated. RBL2 has over 26 reported phosphorylation sites^40–42^, of which hyperphosphorylation is associated with dissociation from E2F4 and RBL2 degradation^40, 43, 44^. Among these phosphorylation sites, phosphorylation of S672 has been shown to be important for attenuating E2F4-RBL2 interactions and is necessary for ubiquitination and degradation^43^. We therefore investigated whether S672 RBL2 phosphorylation status differed between HBECs in ALI and submerged culture. We observed no significant difference in total RBL2 protein expression while S672-phosphorylated RBL2 (S672-RBL2) was significantly higher in HBECs cultures at the ALI compared to submerged (**Fig. 5b-d**). Immunoblotting analysis confirmed that MCC differentiation was activated, evidenced by the induction of Multicilin transcriptional targets, FOXJ1 and TP73, at D4 ALI differentiation (**Fig. 5b**). These data suggest that the RBL2-dependent regulation of Multicilin activity, during ALI differentiation, is associated with increased RBL2 phosphorylation.

Expression of Multicilin in HBECs is known to be dependent on inhibition of NOTCH signaling downstream of air exposure^13^. To determine if RBL2 phosphorylation in differentiating HBECs can be regulated by NOTCH inhibition and Multicilin activity, independent of ALI, we inhibited NOTCH signaling using 5 µM of the γ-secretase inhibitor (DAPT) and induced Multicilin for 4 days in HBECs submerged in differentiation media. Immunoblotting analysis revealed that inhibition of NOTCH signaling alone decreased total RBL2 protein expression, while Multicilin induction alone increased S672-RBL2 expression; both conditions resulted in increases in S672-RBL2:total RBL2 (**Fig. 5e-f)**. Interestingly, only the combination of NOTCH-inhibition and Multicilin-induction significantly increased the S672-RBL2:total RBL2 compared to each factor alone (**Fig. 5f- g**). This data supports a regulation of RBL2 phosphorylation downstream of both NOTCH-inhibition and Multicilin-induction and suggests that they work synergistically to phosphorylate RBL2 at S672.

### RBL2 knockdown increases the transcriptional activity of Multicilin in submerged HBECs

As RBL2 phosphorylation is associated with dissociation from the E2F proteins allowing transcriptional activation, we depleted RBL2 in submerged Multicilin-inducible HBECs to determine whether Multicilin-dependent transcriptional activation of multiciliogenesis would be potentiated independent of regulation by ALI differentiation. Multicilin expression was induced for 4 days in submerged cultures. Induction alone (red bars, **Fig 6a**) had no effect on *RBL2* expression, but significantly increased expression of MCC genes *TP73*, endogenous *MCIDAS*, and *FOXJ1* (**Fig 6a)**. RBL2 knockdown alone had no significant impact on *TP73* gene or protein expression (**Fig. 6a-d**), however, increased MCC gene expression was observed for *FOXJ1* (**Fig 6a**). Interestingly, the combination of RBL2-knockdown and Multicilin-induction led to a significant increase in the expression of MCC genes *SAS6*, *DEUP1*, *TP73* and *FOXJ1* (blue bars + DOX, **Fig. 6a**), the number of TP73-positive cells (**Fig. 6b-c**), and total TP73 protein expression (**Fig. 6d**) when compared to either condition alone. Additionally, in some cells multi-nucleated basal bodies and cilia axonemes were visible, shown in confocal images for pericentrin and ATUB (**Fig. 6e**). We also found that Multicilin-induced MCCs with RBL2 depletion had active ciliary beating, observed in **Supplemental Movie S1**, indicating functional cilia differentiation. These data support a new role for RBL2 in the direct inhibition of the activity of Multicilin. Only in the absence of RBL2 can Multicilin efficiently drive the transcriptional network necessary for multiciliogenesis.

**Figure 6:**
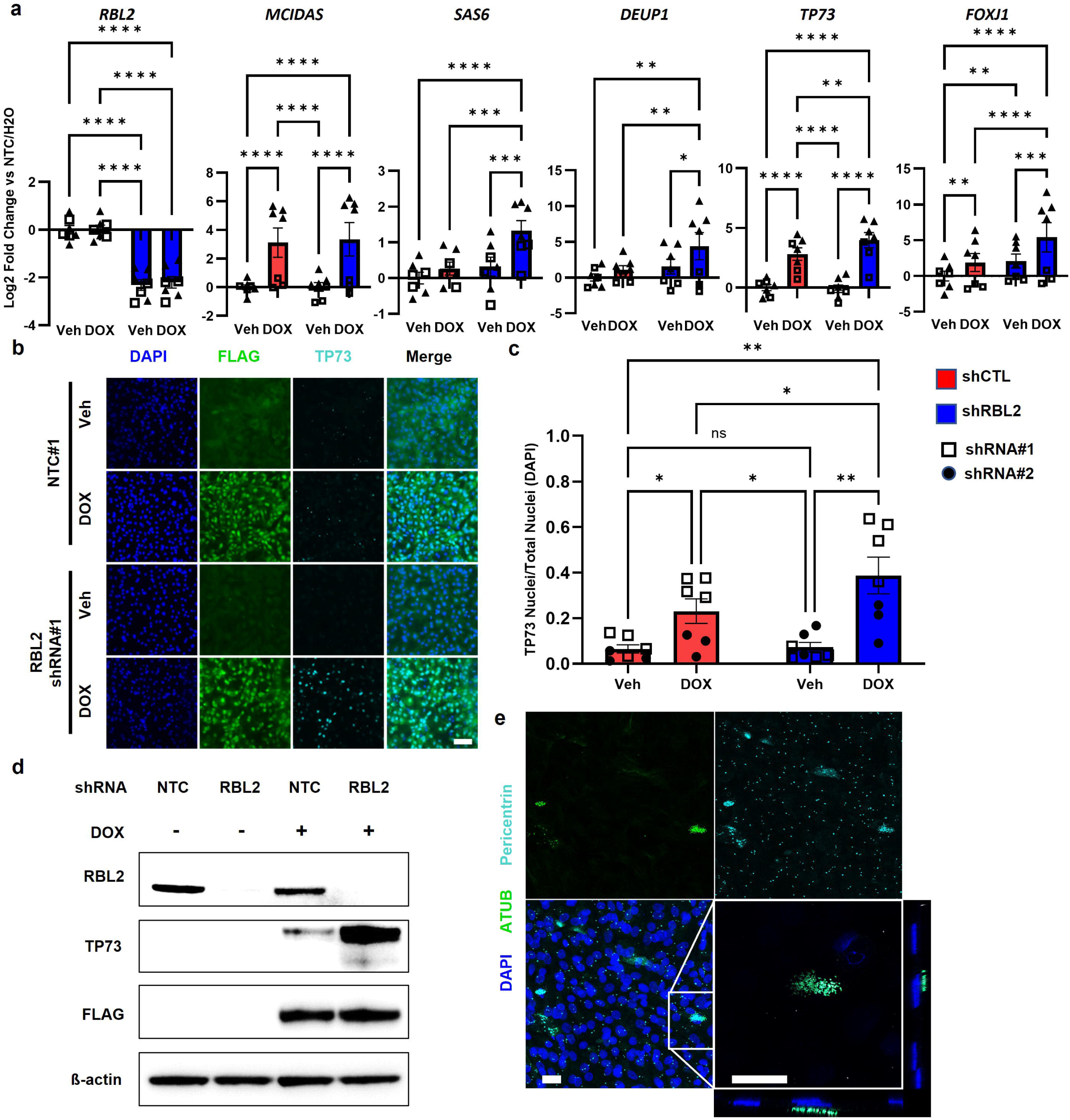
RBL2 inhibits the activity of Multicilin in HBECs. **(a)** RT-qPCR for HBECs comparing non-targeting controls (shNTC#2, red) to RBL2 (shRBL2#2, blue) in the presence (DOX) or absence (Veh, H_2_O) of Multicilin induction. Gene expression is relative to RPLP0 and normalized to the NTC shRNA. N=4 biological replicates. (**b**) Representative IF images for FLAG (Green), TP73 (Cyan) and DNA (DAPI, blue) in HBECs. Scale bars represent 200 µm. (**c**) Quantification of TP73 positive cells. N=3-4 biological replicates each with 2 experimental replicates (**d**) Representative Western blot for RBL2, TP73 and FLAG in HBECs comparing shNTC#2 to shRBL2#2 in the presence (DOX) or absence (Veh) of Multicilin induction (**e**) Representative confocal images of basal bodies (Pericentrin, cyan) and cilia (acetylated α-tubulin, ATUB, green) with DNA counterstain (DAPI, blue) in shRBL2 with Multicilin induction. Scale bars represent 25 µm. Data represents mean±SEM and is compared using a 2-way ANOVA and post-hoc Tukey’s multiple comparisons test (a&c) with significance at *PC<C0.05, **PC<C0.01, ***PC<C0.001, ****PC<C0.0001.

## Discussion

This study was designed to elucidate the mechanisms that regulate the potential of human airway epithelial cells to become multiciliated. We previously reported that Multicilin can drive ectopic multiciliogenesis in MEFs, but only when using a modified version of E2f4 where the C-terminus is replaced with a VP16 transcriptional activator^33^. Here we show that E2f4-VP16 disrupts the interaction of RBL2 with the Multicilin/E2F protein complex, and that depletion of RBL2 alone increases the transcriptional activity of Multicilin in both MEFs and HBECs. Additionally, culture of HBECs at the ALI increases the levels of phosphorylated S672-RBL2 and disrupts the interaction of Multicilin with RBL2. This phenotype was recapitulated in submerged cells only in the presence of both Multicilin-induction and NOTCH-inhibition. Collectively, the data support a novel mechanism whereby the ALI facilitates phosphorylation of RBL2, downstream of the inhibition of NOTCH signaling, allowing the Multicilin-E2f4 transcriptional complex to drive multiciliogenesis independently of cell-cycle mediated RBL2 phosphorylation. This new mechanism is summarized in the schematic in **Figure 7**.

**Figure 7:**
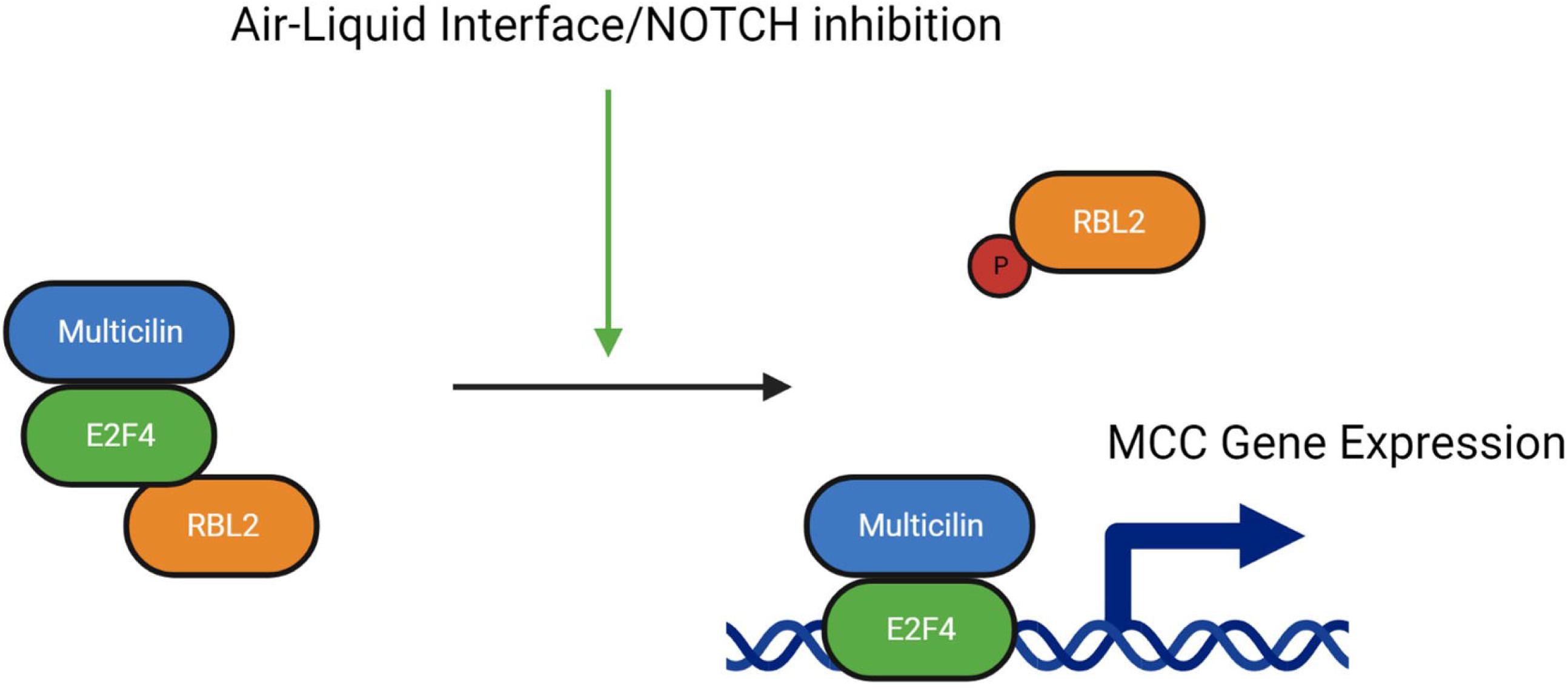
Regulation of Multiciliogenesis and the activity of Multicilin by RBL2. Schematic depicting the new mechanism of regulating MCI activity during multiciliogenesis. Unphosphorylated RBL2 binds to E2F4 and inhibits Multicilin-induction of MCC gene expression. Differentiation at an air-liquid interface and inhibition of NOTCH signaling both phosphorylate RBL2 and dissociate it from the E2f4/Multicilin complex, allowing for induction of MCC gene expression.

Multicilin and GEMC1 initially were associated with the cell cycle as Geminin superfamily proteins through sequence homology of their coiled-coiled domains with Geminin, a negative regulator of DNA replication^24, 45^. Data from a variety species, however, indicates that Multicilin and GEMC1 expression is necessary for proper MCC differentiation^17, 24–27, 45–48^, with loss of function in either gene causing reduced generation of motile cilia^27, 48^, without apparently leading to general cell cycle defects. Both GEMC1 and Multicilin lack a known DNA binding domain, and instead use their conserved C-terminal TIRT domains to interact with the transcription factors E2F4 and E2F5^28, 48^. GEMC1 has been shown to more effectively complex with E2F5 and is thought to specify MCC fate by driving expression of Multicilin and other downstream MCC transcriptional regulators, while Multicilin complexes with both E2F4 and E2F5^28, 48^ to potentiate multiciliogenesis through a positive feedback loop, activation of massive centriole biogenesis, and upregulating the expression of downstream transcription factors. E2F4 and E2F5 have been well characterized as repressors of cell cycle progression, through their interactions with RBL1 and RBL2, which recruit various epigenetic regulators to repress expression of cell cycle genes^49–52^. The mechanism by which Multicilin co-opts E2F4 to induce transcription is largely unexplored, however recent work has shown that the GEMC1 and MCIDAS complex with TP73^53^ and TRRAP^48^ and that their TIRT domains confer differential affinity for SWI/SNF subcomplexes^48^. E2F4 and E2F5 are dual role transcription factors, both acting in essentially all cell types as repressors during the cell cycle, but as activators in progenitors during multiciliogenesis; how the cell differentially regulates the function of E2F4 and E2F5 in this context is still unknown.

Our previously published work demonstrated that multiciliogenesis can be induced in MEFs, a mesenchymal cell type that is not innately capable of MCC differentiation, through a process requiring serum starvation and expression of Multicilin and E2f4ΔCT-VP16^33^. This suggested that a specific cellular state is required to potentiate multiciliogenesis downstream of Multicilin induction, specifically requiring G0/G1 arrest as well as additional unknown factors that can facilitate the role of E2f4 as a transcriptional activator. Here our identification of Rbl2 as a repressor of Multicilin, an interaction which can be interrupted by replacing the PBDB with VP16, is not surprising as serum starvation has been shown to increase hypo-phosphorylation of Rbl2 and E2f-Rbl2 interactions in the MEF derived cell line, NIH3T3^54^. As Rbl2-depleted MEFs expressing FLAG-MCI were able to generate cilia akin to that of Multicilin-induced MEFs expressing E2f4VP16, it is likely that MEFs have an innate capacity to utilize endogenous E2f4 to act as a transcriptional co-activator, however this is reduced by the presence of active Rbl2.

Interestingly, Rbl1 was not efficiently co-immunoprecipitated with FLAG-MCI when expressed alone in MEFs. Multicilin-RBL1 interactions did not appear to be altered by ALI differentiation (data not shown), despite RBL1, and not RBL2, being identified in complex with Multicilin and GEMC1 in recently reported experiments performed in HEK293 cells^48^. RBL2 has been reported to associate with E2F4 during G0 and G1, while RBL1/E2F4 complexes form mainly during S-phase of the cell cycle^55^. This may explain why we detected Rbl1 complexed with Multicilin only when wildtype E2f4 was overexpressed, as endogenous E2f4 would preferentially bind Rbl2, but in the presence of excess E2f4, complexes with E2f4 were able to form. Additionally, our Rbl1 knockdown did not increase MCC specification. Taken together this points to a mechanism where RBL1 regulates GEMC1/E2F5 specification of MCCs, while RBL2 regulates the potential of Multicilin/E2F4 to drive multiciliogenesis.

In conclusion, we provide evidence to support a mechanism by which ALI culture decreases the interaction of RBL2 with Multicilin. Submersion is believed to mimic a hypoxic environment and inhibit differentiation by maintaining NOTCH signaling^13^. We show here that RBL2 interaction with Multicilin is regulated by its phosphorylation and only the combination of NOTCH-inhibition and Multicilin-induction was capable of increasing phospho-RBL2 in submerged cultures, suggesting that RBL2 phosphorylation is downstream of NOTCH-inhibition.

## Materials and Methods

### Cell culture

#### Mouse embryonic fibroblasts (MEFs)

Freshly prepared primary MEFs were obtained from the Genome Manipulation Core at the Salk Institute (strain DR4, non-irradiated). MEFs were grown in DMEM supplemented with 10% FBS and 0.1 mM non-essential amino acid (#11140-050; Invitrogen) in a 37°C humidified incubator with 5% CO2. Only MEFs cultured for less than 4 passages were used in experiments.

#### Human bronchial epithelial cells (HBECs)

Primary HBECs were either a gift from Dr. Steve Brody (Wash U) or isolated from explant donor lungs as previously described^56^ and cultured for one passage in Airway Epithelial Growth Medium (Promocell C-21160). Continued passaging of HBECs was performed in Pneumacult-Ex Plus (StemCell Technologies #05040) and differentiation of cells was performed in differentiation media, either PnuemaCult-ALI (StemCell Technologies #05001) or VALI media^57^ on Transwell inserts (Corning #3470, #3460). Expansion and differentiation of HBECs was performed according to the Pneumacult-ALI manufacturer guidelines, with the exception that all cell culture surfaces were coated in PureCol (Advanced Biomatrix #5005) according to product guidelines. The use of de-identified HBECs was approved by the IRB of both the University of Southern California and the University of Iowa as “not Human Subjects Research.”

### Virus production and infection

#### pAd/CMV/V5 expressing mouse Multicilin and E2f4 proteins

The plasmids used in this study are modified from those described in Kim *et al*^33^. Mouse Multicilin (GenBank: AK134107.1) was obtained from the Riken mouse FANTOM clone library (clone ID 5830438C23). Mouse E2f4 (NCBI Ref: NM_148952.1) was derived from a cDNA obtained from Dharmacon (clone ID 4987691), by using the full-length coding domain (wildtype) or by using a C-terminal truncation (1–774bp) that was replaced with the transcriptional activation domain of VP16 (Viral Protein 16; amino acids 413–490) from the UL48 gene of Herpes Simplex Virus-1 (HSV-1; GenBank: KM222726.1). The NLS based on SV40 T-antigen (PKKKRKV) and the 6xmyc tags were derived from the CS2 vectors. S2 cleavage sequence used in this study was T2A. Genes assembled in a pENTR/D-TOPO vector (K240020, Invitrogen) were validated by sequencing, and then transferred to the pAd/CMV/V5-DEST vector (#V49320; Invitrogen) using Gateway cloning.

#### Adenovirus

Adenoviruses were generated by GT3 core at the Salk Institute. Vector DNAs were initially transfected into 293T cells, validated for intact protein expression by immunoblot and immunostaining, and then used to generate crude adenovirus lysates and titer calculated. Primary MEFs were plated with 10% FBS-DMEM for overnight, then treated with 2% FBS-DMEM for 1 day before adenovirus infection. Adenovirus infection was performed by adding the adenovirus crude lysate to the MEF cells for four hours, washing with pre-warmed PBS once, then growing infected cells with 2% FBS-DMEM. The cells were subjected to relevant analysis according to days post-infection (PI).

#### Lentivirus production and infection

Lentiviruses were generated in either the GT3 Core Facility of the Salk Institute or University of Iowa Viral Vector Core (http://www.medicine.uiowa.edu/vectorcore). Additionally, lentivirus was also prepared using the calcium-phosphate transfection method^58^ and third generation lentiviral packaging plasmids, pMDLg/RRE, pRSV-Rev, and pMD2.G (#12253, #12251, #12259; Addgene) as previously described^59^. RBL2 and non-targeting control shRNA lentiviral transfer plasmids were either purchased from Sigma or generated by cloning sequences into pSicoR-Ef1a-mCh-Puro-Puro (#31847;Addgene, see **Supplemental Tables S2&S3**) as previously described^60^. Doxycycline inducible-FLAG-Multicilin lentiviral transfer plasmid was received as a gift from Alvarez R & Verma IM, (Unpublished).

### Gene Knockdown (KD) Experiments

#### RBL2 KD via siRNA transfection

MEFs were grown in 10% FBS-DMEM overnight, then treated with 2% FBS-DMEM. Six hours after serum-reduced treatment, siRNAs were transfected into MEFs using Lipofectamine RNAiMAX (Invitrogen, #13778150) according to the manufacturer’s instructions. The transfection is performed by adding the pre-mixed complex of siRNA with the reagent in reduced serum media (Opti-MEM; Invitrogen, #31985062) to cells without a media change. Eighteen hours later, the cells were washed with 2% FBS-DMEM and infected with the indicated adenovirus. Four hours after infection, the cells were washed with pre-warmed PBS once, 2% FBS-DMEM once then grown under serum-reduced condition with 2% FBS-DMEM. The cells were subjected to further analysis according to days post-infection (PI). The siRNA for negative control (#12935–113) and siRNAs against Rbl1 (AM16708 ID#151420), Rbl2#1 (AM16708 ID#151423), Rbl2#2 (AM16708 ID#68825), Rbl2#3 (AM16708 ID#68921) were purchased from Thermo Fisher Scientific. The oligo sequences of siRNAs used in this study; siRbl1 (5′-GCU AAG UUA AGC UUA AUA Ctt-3′), siRbl2#1 (5′-CCU UCA UUG GUU AGC AUG Utt-3′) and siRbl2#2 (5′-GGG AAA UGA CCU UCA UUG Gtt-3′), siRbl2#3 (5’-GGG ACC GCU GAA GGA AAC Utt-3’).

#### RBL2 knockdown via shRNA transduction

HBECs cultured in Pneumacult-Ex Plus were dissociated into single cell suspensions using Accutase (#AT 104, Innovative Cell Technologies), counted, and the appropriate number of cells per infection condition were pelleted by centrifugation at 400g for 5 minutes. Cells were then resuspended at a density of 500K cells/mL in lentivirus diluted to an MOI of 20 in Pneumacult-Ex Plus with 10 µg/mL polybrene (#TR-1003, Millipore Sigma) and incubated at 37°C for 2 hours, vortexed every 20 minutes to prevent cell clumping. Without removing the virus, cells were seeded at the appropriate cell density for either cell expansion or air-liquid interface differentiation and then incubated overnight 37°C. Virus was removed the next day and cells were cultured in 1µg/mL puromycin for at least three days before use in experiments.

### Real time quantitative polymerase chain reaction (qRT-PCR)

Total RNA was isolated from experimental MEFs (Quick-RNA prep kit; ZYMO RESEARCH, #R1054) or HBECs (Quick-RNA Microprep Kit; ZYMO RESEARCH, #R1051) and at least 500ng RNA was converted in cDNA using High-Capacity cDNA Reverse Transcription Kit (Applied Biosystems, # 4368814). Gene expression was then assayed by quantitative PCR using PowerUp SYBR (Applied Biosystems, #A25742) and appropriate primer pairs in triplicate on the ABI Prism 7900HT thermal cycler (Applied Biosystems, #4329001) or the QuantStudio5 (Applied Biosystems, # A28575) . Gene expression in HBECs and MEFs was normalized using reference genes *RPLP0* and *Gapdh* respectively and calculated as log2 fold change versus the average of control samples as stated. Primer sequences are listed in **Supplemental Table S2**.

### Immunoprecipitation, immunoblotting analysis, and silver staining

For identifying proteins interacting with FLAG-tagged Multicilin by MASS analysis, MEFs were infected by pAd/CMV/V5 vector encoding 3xFLAG-Multicilin, NLS-6xmyc-E2f4ΔCt-VP16-T2A-Multicilin, and NLS-6xmyc-E2f4WT-T2A-Multicilin. Non-infected MEFs were used as the negative control. Two days after infection, the non-infected MEFs or infected MEFs were lysed in lysis buffer (50mM HEPES-KOH [pH 7.5], 500NaCl, 5mM EDTA [pH 8.0]), 2% Triton, DNase I (10ug/ml)) by incubation on ice for 20min. Lysates were cleared by centrifugation at 12,000 rpm for 20 min at 4 °C, then precleared by incubation with Protein G agarose (Invitrogen #20397) agitating for 1hr at 4°C. Spun-down beads were incubated with FLAG M2 agarose (Sigma #A2220) agitating for 2.5 hours at 4°C. The beads were washed twice with wash buffer (1:1 mix of 50mM

HEPES and Lysis buffer), washed with lysis buffer twice, then washed twice again with wash buffer. Spun-down beads were incubated with 40uL of FLAG peptide (500ng/mL) for bound protein elution overnight at 4°C. Elutes were subjected to protein sampling for further immunoblot analysis and silver staining. Eluted protein samples and experimental protein lysates were subjected to SDS-PAGE and transferred to a PVDF membrane that was then blocked with 0.5% casein blocker (#161-0783, Bio-Rad) in PBS for 30 min followed by incubation with the indicated primary antibodies overnight at 4 °C in 0.1% Tween-20 in blocking buffer. After extensive washing in 0.1% Tween-20 in PBS, the blots were incubated with Alexa 680 or 800-conjugated anti-mouse or rabbit secondary antibodies (Invitrogen, A21058, A21076, A32735) for 45 min at room temperature, washed with 0.1% Tween-20 in PBS, and imaged using Odyssey (LI-COR). Silver staining was performed with Pierce Silver Stain Kit (Thermo #24612) based on the manufacturer’s instructions.

### Mass Spectrometry analysis

Samples were precipitated by methanol/ chloroform and redissolved in 8 M urea/100 mM TEAB, pH 8.5. Proteins were reduced with 5 mM tris(2-carboxyethyl)phosphine hydrochloride (TCEP, Sigma-Aldrich) and alkylated with 10 mM chloroacetamide (Sigma-Aldrich). Proteins were digested overnight at 37°C in 2 M urea/100 mM TEAB, pH 8.5, with trypsin (Promega). Digestion was quenched with formic acid, 5 % final concentration. The digested samples were analyzed on a Fusion Orbitrap tribrid mass spectrometer (Thermo). The digest was injected directly onto a 30 cm, 75 um ID column packed with BEH 1.7um C18 resin (Waters). Samples were separated at a flow rate of 300 nl/min on a nLC 1000 (Thermo). Buffer A and B were 0.1% formic acid in water and 0.1% formic acid in 90% acetonitrile, respectively. A gradient of 1-35% B over 110 min, an increase to 50% B over 10 min, an increase to 90% B over 10 min and held at 90%B for a final 10 min was used for 140 min total run time. Column was re-equilibrated with 15 µl of buffer A prior to the injection of sample. Peptides were eluted directly from the tip of the column and nanosprayed directly into the mass spectrometer by application of 2.5 kV voltage at the back of the column. The Orbitrap Fusion was operated in a data dependent mode. Full MS scans were collected in the Orbitrap at 120K resolution with a mass range of 400 to 1600 m/z and an AGC target of 5e5. The cycle time was set to 3 sec, and within these 3 secs the most abundant ions per scan were selected for CID MS/MS in the ion trap with an AGC target of 1e4 and minimum intensity of 5000. Maximum fill times were set to 50 ms and 100 ms for MS and MS/MS scans, respectively. Quadrupole isolation at 1.6 m/z was used, monoisotopic precursor selection was enabled, and dynamic exclusion was used with exclusion duration of 5 sec. Protein and peptide identification were done with Integrated Proteomics Pipeline – IP2 (Integrated Proteomics Applications). Tandem mass spectra were extracted from raw files using RawConverter^61^ and searched with ProLuCID^62^ against Uniprot mouse database. The search space included all fully-tryptic and half-tryptic peptide candidates. Carbamidomethylation on cysteine was considered as a static modification. Data was searched with 50 ppm precursor ion tolerance and 600 ppm fragment ion tolerance. Identified proteins were filtered to using DTASelect^63^ and utilizing a target-decoy database search strategy to control the false discovery rate to 1% at the protein level^64^.

### Immunocytochemistry and image processing

MEF centriolar and cilia imaging was performed as previously described^33^. HBECs were grown on Corning 6.5mm Transwell inserts (#3470) or Nunc Lab-Tek chamber slides (#177445) and fixed in 4% paraformaldehyde in PBS overnight at 4°C. Cells were concurrently permeabilized and blocked-in antibody dilution buffer (5% bovine serum albumin, 0.3% Triton X-100, PBS) before incubation with antibodies (**Supplemental Table S4**) in antibody dilution buffer. Samples were incubated at room temperature with primary antibodies for 2 hours and secondary antibodies for 45 minutes. Counterstains in PBS were performed directly after secondary antibody incubation (DAPI 1µg/mL 5 minutes, Phalloidin (Biolegend#424203) 5 units/mL), with (3x) 5-minute PBS washes performed after primary antibody and counterstain incubations. Samples were mounted with No. 1.5 coverslips using Fluoromount-G mounting medium (Invitrogen #00-4958-02). Fluorescent image tile scans were performed on either the DMi8 (Leica) or Revolution (Echo) fluorescent microscope systems and processed in Fiji^65^. TP73 positive cells were quantified by thresholding for positive DAPI and TP73 signal based on control samples, water shedding and then analyzing particle counts to give the number of nuclei positive for each stain. Cilia area coverage was calculated by thresholding for positive acetylated α-tubulin staining and then dividing by the area covered by counterstain (Phalloidin), with a threshold set high enough to give positive signal for the entire cell coverage area. For Cilia quantification three independent experiments were performed in each of which 153-310 cells were counted per experiment totaling 676 cells for siCTL, 660 cells for siRbl2 and 651 cells for E2f4VP16.

### RNA sequencing

Total RNA was isolated using Quick-RNA prep kit (Zymo Research Cat#R1054). RNAseq libraries were constructed with Illumina Truseq RNA Sample Preparation kit v2 according to the manufacturer’s instructions and sequenced on a HiSeq 2500 at 1x50 base pairs to a depth of 20-40 million reads. Each RNAseq condition was performed in triplicate with RNAs isolated from three individual experiments using MEFs. Preprocessing of fastq files and differential expression analysis was performed using tools available on Galaxy server (https://usegalaxy.org/). Quality check and adapter trimming of the reads were performed with fastQC (http://www.bioinformatics.babraham.ac.uk/projects/fastqc) and FastP tools^66^ respectively. Reads were mapped to the mouse reference genome assembly (GCF_000001635.27_GRCm39_genomic.fna) and BAM files were generated using RNA-STAR^67^. Further genomic feature was assigned with featureCounts^68^ using mouse genomic coordinates file (GCF_000001635.27_GRCm39_genomic.gtf). Finally differential gene expression analysis was performed using DEseq2 tool^69^. For differential expression studies, genes with Log2 fold change ≥ 2 (compared to Control/NTC) were considered and adjusted p value ≤ 0.05 was set as threshold value for statistical significance. r-Log normalized counts for genes with significant differential expression were used to generate heatmaps and Euclidean clustering method was applied. pHeatmap, ggplot2, dplyr, tidyverse and their dependency packages were used to generate heatmaps and volcano plots in RStudio (RStudio Team, 2020; R Core Team, 2021). Codes and processed data files used for volcano plots and Heatmaps is available in Git-Hub repository (https://github.com/gautam-lk/RyanLab_RBL2). Scatterplots were prepared in GraphPad Prism (Version 9.4.1) and a web-based tool, InteractiVenn, (http://www.interactivenn.net/index2.html) was used to generate Venn diagram(s). The Gene Ontology based biological, cellular and molecular functions were predicted using g:Profiler (https://biit.cs.ut.ee/gprofiler/gost) (**Supplemental Table S5**).

### Cilia beat frequency analysis

ALI inserts were placed on glass bottom dishes (VWR #75779-574) and cilia movement was captured using a 60x water immersion objective on an inverted Zeiss microscope system equipped with a heated CO_2_ chamber and highspeed camera with video capture and CBF analysis performed using Sisson-Ammons Video Analysis Software (SAVA). At least 5 random field of views with beating cilia were captured per sample, with at least 3 samples analyzed per condition.

### Statistical Analysis

Data was analyzed in GraphPAD Prism 9.4.1 and are expressed as the mean with error bars representing SEM. Data represents 4-6 biological replicates and 1-3 experimental replicates or two independent sets of target and control shRNA, per experiment, as indicated in the figure legends. Paired biological samples are denoted in each figure by data point symbol shape when not otherwise labeled. Comparison of treatment to control groups was performed by two-tailed students paired t-test or ratio paired t-test, as appropriate. Comparisons of multiple groups was performed by 2-way ANOVA followed by post-hoc Tukey’s multiple comparison test. P value significance is as follows: *PC<C0.05, **PC<C0.01, ***PC<C0.001, ****PC<C0.0001.

## Acknowledgements

This project was funded by NIH NHLBI HL139828 (ALR, CK, ZB) and HL153622 (ALR) and supported by the Mass Spectrometry Core and the GT3 Core Facility of the Salk Institute with funding from NIH-NCI CCSG: P30 014195, an NINDS R24 Core Grant, the Helmsley Center for Genomic Medicine and funding from NEI. We thank J. Moresco, J. Diedrich for technical support.

## Author contributions

Conceptualization, A.L.R. and C.K.; Methodology, E.J.Q., S.K. C.K. and A.L.R.; Investigation, E.J.Q., S.K.; Formal Analysis, E.J.Q., S.K., L.K.G.; Writing – Original Draft, E.J.Q. and A.L.R.; Writing – Review & Editing, E.J.Q., S.K., Z.B., C.K. and A.L.R.; Funding Acquisition, A.L.R.; Supervision, A.L.R, Z.B. and C.K.

## Data Availability

The RNA sequencing data generated in this study has been deposited in the NCBI Gene Expression Omnibus (GEO) (https://www.ncbi.nlm.nih.gov/geo/) under accession code GSE235682. The main data supporting the findings of this study are available within the article and its Supplementary Figures and Supplementary Tables.

## Code Availability

The customized code used for RNAseq data analysis can be found at https://github.com/gautam-lk/RyanLab_RBL2.

