## Supplemental Information for "RBL2 represses the transcriptional activity of Multicilin to inhibit multiciliogenesis"

**Title**

**Authors and Affiliations**

Erik J. Quiroz^1,2,3*^, Seongjae Kim^4,5*^, Lalit K. Gautam^1^, Zea Borok^5^, Christopher Kintner^4^, Amy L. Ryan^1,2,3*^

^1^Department of Anatomy and Cell Biology, Carver College of Medicine, University of Iowa, Iowa City, IA 52240

^2^Hastings Center for Pulmonary Research, Division of Pulmonary, Critical Care and Sleep Medicine, Department of Medicine, University of Southern California, Los Angeles, CA 90033

^3^Department of Stem Cell Biology and Regenerative Medicine, University of Southern California, Los Angeles, CA 90033

^4^The Salk Institute of Biological Studies, La Jolla, CA 92093

^5^Division of Pulmonary, Critical Care and Sleep Medicine, Department of Medicine, University of California San Diego, San Diego, CA 92037

*These authors contributed equally to this manuscript

**Corresponding Author** **Contact Information**

Amy L. Ryan, PhD

Associate Professor: Anatomy and Cell Biology
Associate Director: Center for Gene Therapy

BSB, 1-400 Core
University of Iowa
51 Newton Road
Iowa City, Iowa, 52242


**Supplementary Information**

The raw data supporting the data provided in the manuscript can be accessed on figshare. Raw data files associated with: Figure 1 <https://doi.org/10.6084/m9.figshare.23792361.v1>. Figure 2 <https://doi.org/10.6084/m9.figshare.23792364.v1>, Figure 4 <https://doi.org/10.6084/m9.figshare.23792181.v1>, Figure 5 <https://doi.org/10.6084/m9.figshare.23792301.v2> and Figure 6 <https://doi.org/10.6084/m9.figshare.23795949>.

**Supplemental Tables**

**Supplementary Table S1: Mass spectrometry analysis of MEFs expression Multicilin and Multicilin/E2f4VP16.** Provided as an excel file

**Supplementary Table S2: Oligonucleotide sequences for RT-qPCR primers**

| **Mouse RT-qPCR Primers** | | |
| --- | --- | --- |
| **Primer Target** | | **Sequence** |
| mRbl2 | Forward | 5’-GAGAAGCTGGAGCGGATACT-3’ |
| mRbl2 | Reverse | 5’-TCTGGCTGGAAATGCTGAGA-3’ |
| mCcno | Forward | 5’-CTGCAGCTCCCTACTCAACC-3’ |
| mCcno | Reverse | 5’-CCTTTCGGAAGTCGTAGCAG-3’ |
| mDeup1 | Forward | 5’-ATATGAGAACGAAAGACTCCGA-3’ |
| mDeup1 | Reverse | 5’-TGGCTAATTCTGTCTCCACTG-3’ |
| mMulticilin | Forward | 5’-CCGAGCCCTTCCAGATCAAG-3’ |
| mMulticilin | Reverse | 5’-TTAGGGTCACGATTGTGCAGG-3’ |
| mGapdh | Forward | 5'-AGGTCGGTGTGAACGGATTTG-3' |
| mGapdh | Reverse | 5'-TGTAGACCATGTAGTTGAGGTCA-3' |
| **Mouse siRNA Sequences** | | |
| **siRNA** | **Catalog#** | **Sequence** |
| siRbl1 | AM16708 ID#151420 | GCU AAG UUA AGC UUA AUA Ctt |
| siRbl2#1 | AM16708 ID#151423 | CCU UCA UUG GUU AGC AUG Utt |
| siRbl2#2 | AM16708 ID#68825 | GGG AAA UGA CCU UCA UUG Gtt |
| siRbl2#3 | AM16708 ID#68921 | GGG ACC GCU GAA GGA AAC Utt |
| siCTRL | #12935–113 | Not Provided |
| **Human RT-qPCR Primers** | | |
| **Primer Target** | | **Sequence** |
| hRBL2 | Forward | 5’-CACCTTGCCAGTTCCACAGC-3’ |
| hRBL2 | Reverse | 5’-TGGAGGAGCATCCATATTTGCCT-3’ |
| hSAS6 | Forward | 5’-GCCTGCACATTCCAGCAGCA-3’ |
| hSAS6 | Reverse | 5’-GTTCCTGAACCAGGGTGGCT-3’ |
| hDEUP1 | Forward | 5’-TGACATGGAGAACCAAGCCCA-3’ |
| hDEUP1 | Reverse | 5’-CGTGTCTCCAAGCCCGCAT-3’ |
| hTP73 | Forward | 5’-GCAAGCGTGCCTTCAAGCAG-3’ |
| hTP73 | Reverse | 5’-CGTGTCCTCGTCTCCATGCC-3’ |
| hFOXJ1 | Forward | 5’-GCTACTTCCGCCACGCAGAT-3’ |
| hFOXJ1 | Reverse | 5’-TTCGTCCTTCTCCCGAGGCA-3’ |
| hCC10 | Forward | 5’-ACCATGAAACTCGCTGTC-3’ |
| hCC10 | Reverse | 5’-TCATAACTGGAGGGTGTGTCC-3’ |
| hMUC5AC | Forward | 5’-ACCAATGCTCTGTATCCTTCCC-3’ |
| hMUC5AC | Reverse | 5’-TGGTGGACGGACAGTCACT-3’ |
| hMCIDAS | Forward | 5’-TGGCGGACCAGAACCAGAGA-3’ |
| hMCIDAS | Reverse | 5’-GTTCGGCTGGCGAGTTCCTT-3’ |
| hRPLP0 | Forward | 5’-CCGTGATGCCCAGGGAAGAC-3’ |
| hRPLP0 | Reverse | 5’-GCATCTGCTTGGAGCCCACA-3’ |

**Supplementary Table S3: Oligonucleotide sequences for si/shRNA**

| **human shRNA Sequences** | | | | |
| --- | --- | --- | --- | --- |
| shRNA | Lentivirus Transfer Plasmid Backbone | Catalog# | Target Sequence | Hairpin Loop |
| NTC#1 | pSicoR-Ef1a-mCh-Puro | N.A. (Cloned from Adggene #31845) | ATATGTCGCCGACCTAGCGAT | TTGGATCCA |
| NTC#2 | pLKO.1 | Addgene #136035 | CCTAAGGTTAAGTCGCCCTCG | CTCGAG |
| RBL2#1 | pSicoR-Ef1a-mCh-Puro | N.A.(Cloned from Adggene #31845) | GCACTTTCTACAATGTGAA | TTCAAGAGA |
| RBL2#2 | pLKO.1 | Sigma Mission RNAi #TRCN0000039923 | GCTGAGAGAAATATGGAACTT | CTCGAG |

**Supplementary Table S4: Antibody Information**

| **Antibody Target** | **Vendor** | **Catalog Number** |
| --- | --- | --- |
| Rbl1 | Proteintech | 13354-1-AP |
| Rbl2 | CellSignaling Technologies | 13610 |
| Stil | Bethyl | A302-441 |
| γ-tubulin | Santacruz | sc-17787) |
| Cep164 | Santacruz | sc-240226), |
| Arl13b | NeuroMab | 75-287 |
| Deup1 | Atlas antibodies | HPA010986 |
| Sas6 | Santacruz | sc-81431 |
| FOXJ1 | eBioscience | 14-9965 |
| RBL2 S672 | abcam | ab284755 |
| FLAG | Sigma | F4042 |
| CCDC39 | Atlas antibodies | HPA035364 |
| RSPH9 | Atlas antibodies | HPA031703 |
| TP73 | Atlas antibodies | HPA027314 |
| ß-actin | Genetex | GT5512 |

**Supplementary Table S5: Metadata for RNAseq analysis of non-infected, Multicilin, Multicilin/siRBL2 and Multicilin/E2f4VP16 MEFs.** Provided as an excel file

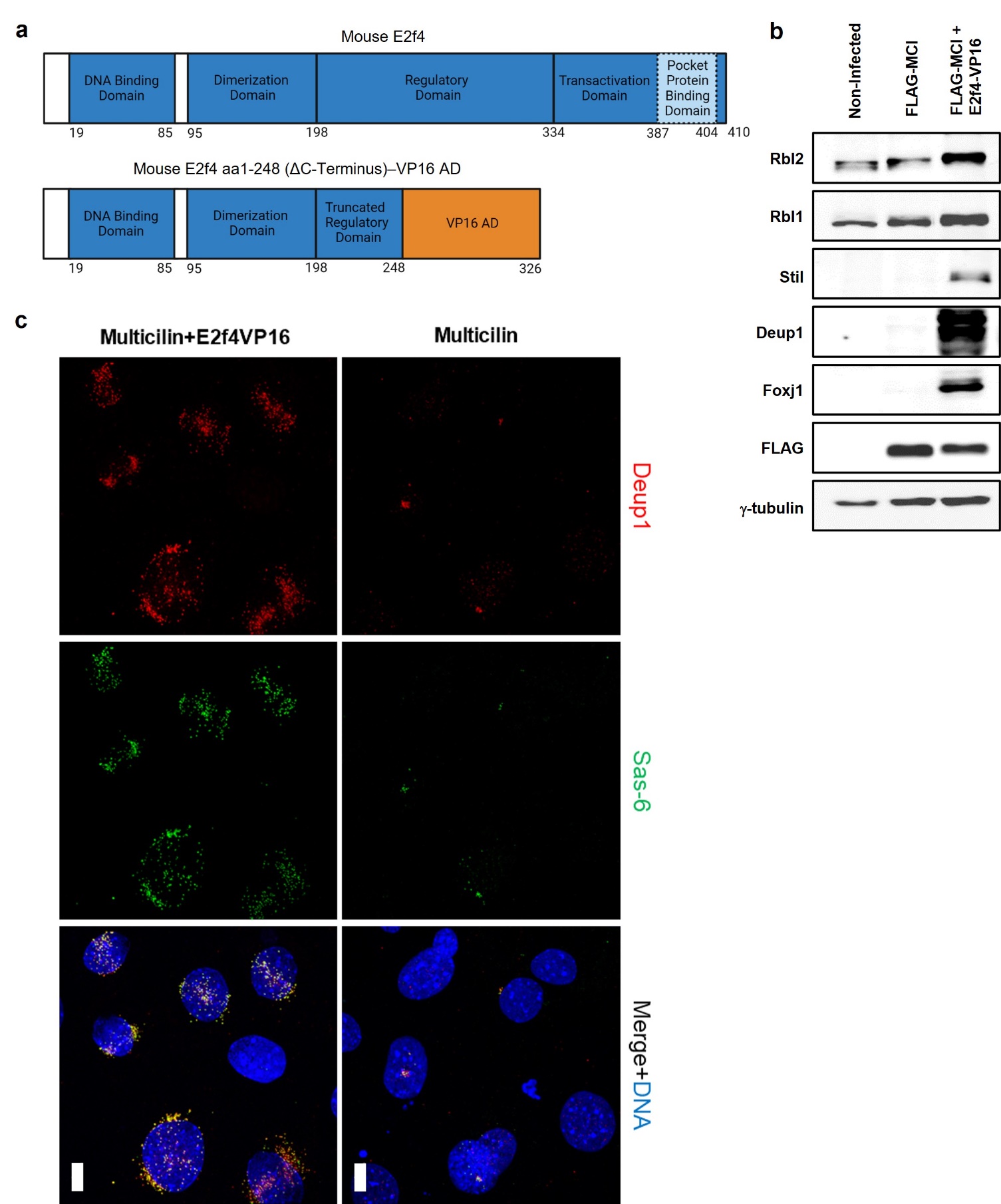
**Supplemental Figures**

**Supplementary Figure S1: MCC protein expression is induced by Multicilin/E2F4-VP16 transduction in MEFs.** (**a**) Diagram depicting mouse E2f4 protein domains and replacement of c-terminal amino acids (aa) 249-410 with VP16 transcriptional activation domain (VP16 AD) (**b**) Western blot comparing non-infected MEFs and MEFs infected with FLAG-MCI or FLAG-MCI+E2f4VP16. (**c**) Confocal imaging of MEFs infected with FLAG-MCI or FLAG-MCI+E2f4VP16 for deuterosome mediated centriolar biogenesis proteins Deup1 (red) and Sas6 (green) with DNA counterstain (DAPI, blue). Scale Bars = 10 µm.

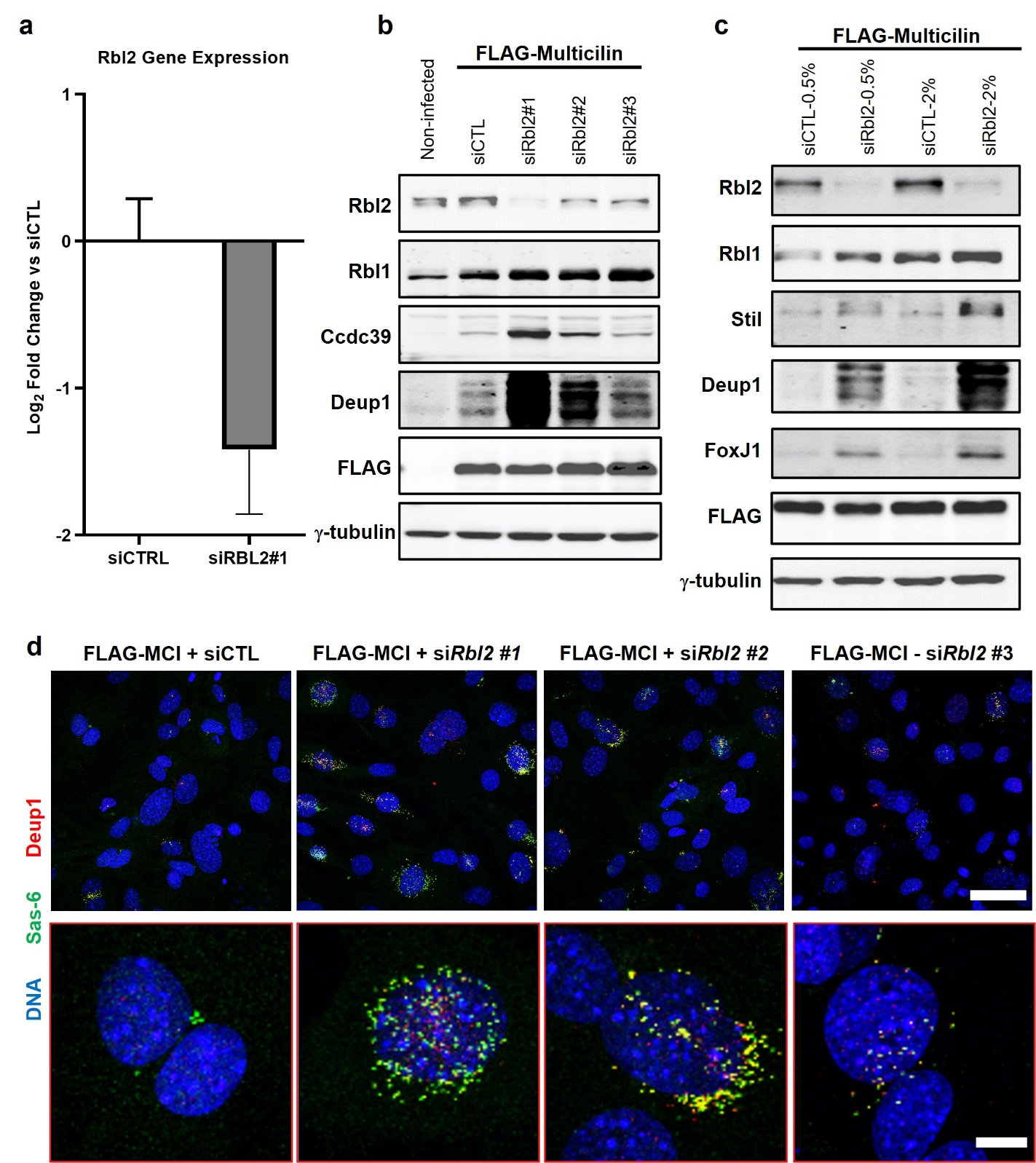

**Supplementary Figure S2: MCC protein expression induced by Multicilin transduction/Rbl2 knockdown in MEFs.** (**a**) qRT-PCR analysis for Rbl2 expression in MEFs in the presence of siRNA targeting Rbl2 (siRBL2#1, siRBL#2) or two non-targeting control siRNA (siCTL#1, siCTL#2). N=2 experimental replicates and 2 technical replicates for each. Gene expression relative to Gapdh and normalized to siCTLs. Data represents mean±SEM. (**b-c**) Western blot analysis with antibodies against given proteins for non-infected MEFs and MEFs infected with FLAG-MCI and transfected with given siRNAs and (**c**) siCTL and siRBL2 at 0.5% and 2% of transfection mix. (**d)** Confocal imaging comparing non-infected to FLAG-MCI ± siRNA-RBL2 for deuterosome mediated centriolar biogenesis proteins Deup1 (red) and Sas6 (green) with DNA counterstain (DAPI, blue). Scale Bars = 50 µm and 10 µm.

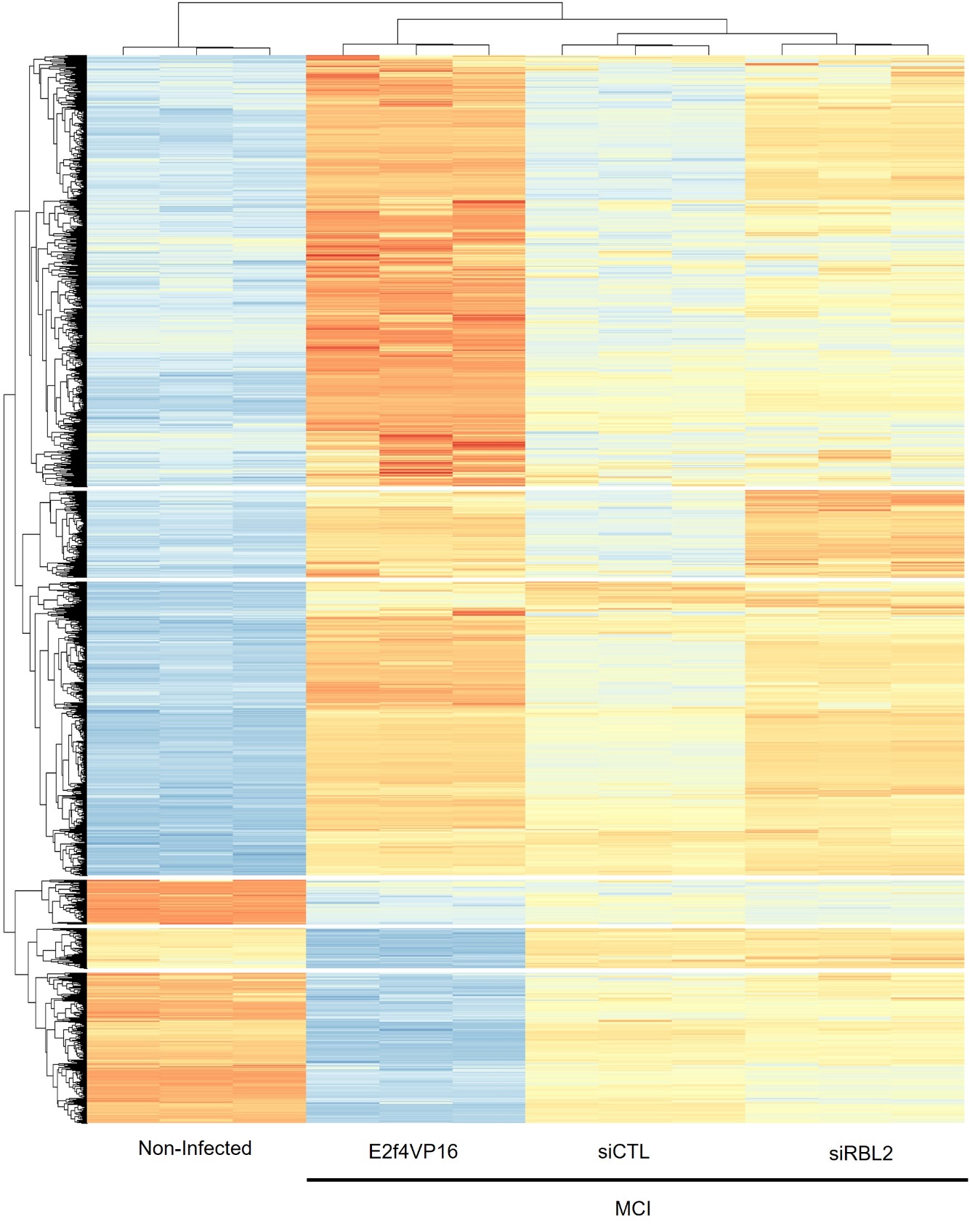
**Supplementary Figure S3: Heatmap of total gene expression comparing Multicilin/siRBL2 and Multicilin/E2f4VP16 MEFs.** Heatmap representing gene expression from RNAseq of non-infected MEFs and MEFs infected with FLAG-MCI+siCTL, FLAG-MCI+siRBL2 or FLAG-MCI+E2f4VP16.

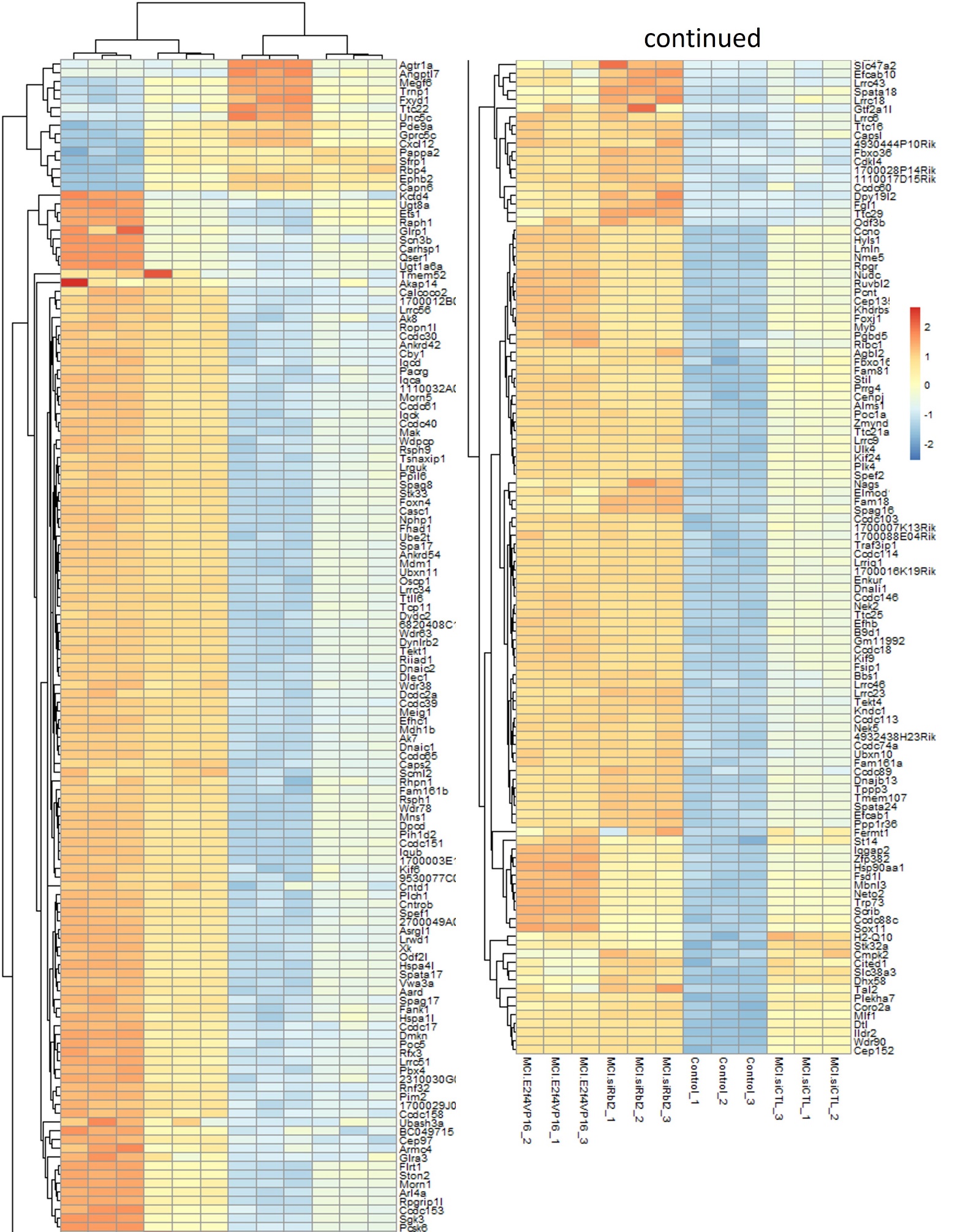
**Supplementary Figure S4: Heatmap of MCC associated gene expression comparing Multicilin/siRBL2 and Multicilin/E2f4VP16 MEFs.** Heatmap representing only MCC-associated gene expression from RNAseq of non-infected compared to FLAG-MCI+siCTL, FLAG-MCI+siRBL2 or FLAG-MCI+E2f4VP16 MEFs.

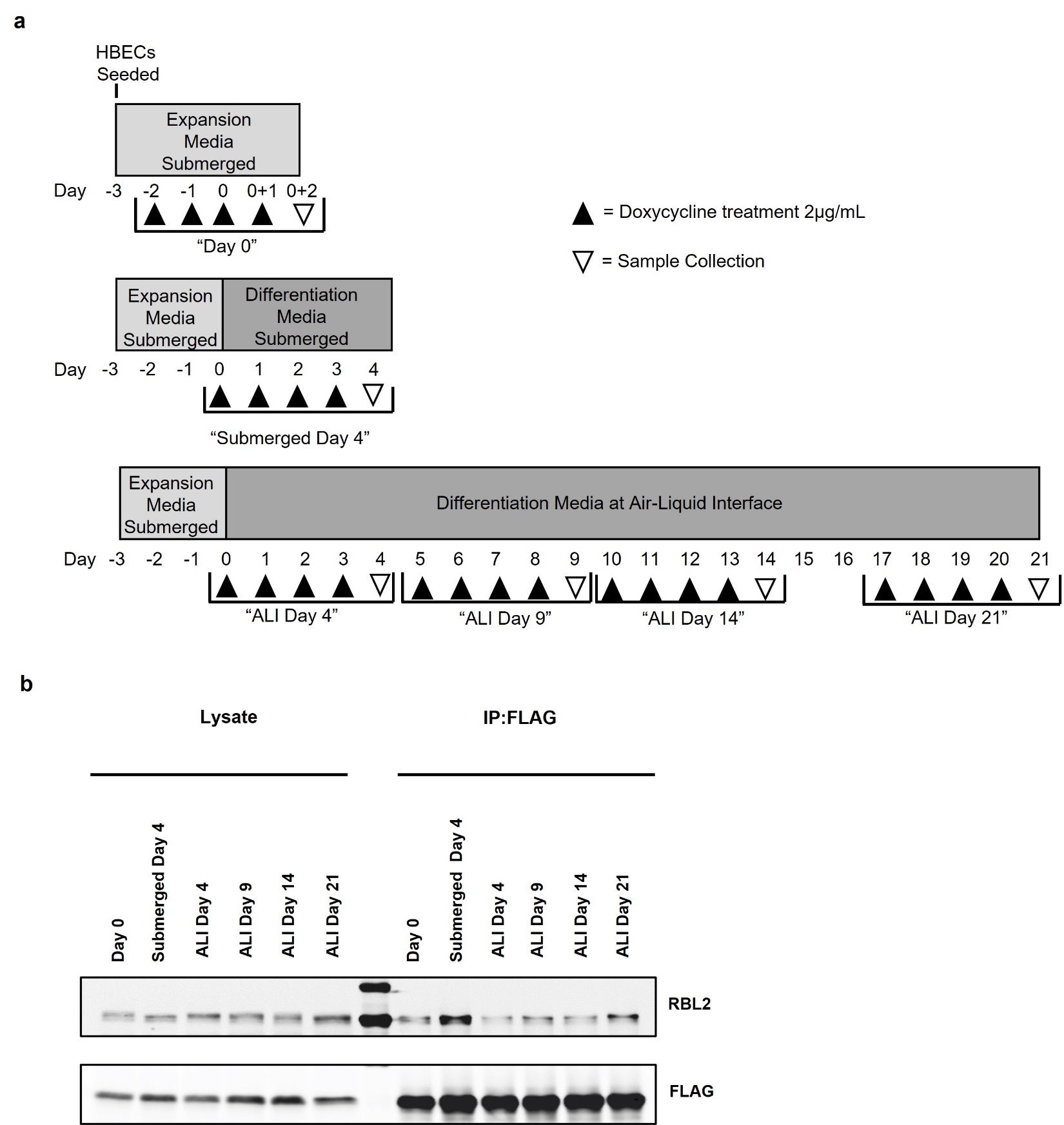

**Supplementary Figure S5: Co-IP of Multicilin and RBL2 in HBECs at the ALI. (a)** Experimental outline for Multicilin induction at the air-liquid interface. **(b)** Western blot analysis for RBL2 and FLAG of total protein and co-IP with FLAG from HBECs transduced with doxycycline inducible FLAG-Multicilin. Cells treated with doxycycline for 4 days prior to sample collection at different given timepoints of air-liquid differentiation (ALI) or after 4 days submerged in differentiation media (Submerged Day 4).

**Supplemental Movie S1: Multiciliated cell cilia beating in submerged HBECs with RBL2 knockdown and Multicilin overexpression**
